# Multilevel irreversibility reveals higher-order organisation of non-equilibrium interactions in human brain dynamics

**DOI:** 10.1101/2024.05.02.592195

**Authors:** Ramón Nartallo-Kaluarachchi, Leonardo Bonetti, Gemma Fernández-Rubio, Peter Vuust, Gustavo Deco, Morten L. Kringelbach, Renaud Lambiotte, Alain Goriely

## Abstract

Information processing in the human brain can be modelled as a complex dynamical system operating out of equilibrium with multiple regions interacting nonlinearly. Yet, despite extensive study of the global level of non-equilibrium in the brain, quantifying the irreversibility of interactions among brain regions at multiple levels remains an unresolved challenge. Here, we present the Directed Multiplex Visibility Graph Irreversibility framework, a method for analysing neural recordings using network analysis of time-series. Our approach constructs directed multi-layer graphs from multivariate time-series where information about irreversibility can be decoded from the marginal degree distributions across the layers, which each represents a variable. This framework is able to quantify the irreversibility of every interaction in the complex system. Applying the method to magnetoencephalography recordings during a long-term memory recognition task, we quantify the multivariate irreversibility of interactions between brain regions and identify the combinations of regions which showed higher levels of non-equilibrium in their interactions. For individual regions, we find higher irreversibility in cognitive versus sensorial brain regions whilst for pairs, strong relationships are uncovered between cognitive and sensorial pairs in the same hemisphere. For triplets and quadruplets, the most non-equilibrium interactions are between cognitive-sensorial pairs alongside medial regions. Finally, for quintuplets, our analysis finds higher irreversibility when the prefrontal cortex is included in the interaction. Combining these results, we show that multilevel irreversibility offers unique insights into the higher-order, hierarchical organisation of neural dynamics and presents a new perspective on the analysis of brain network dynamics.

## INTRODUCTION

The human brain produces complex spatiotemporal neural dynamics across multiple time and length scales. Abstracting the brain as a large-scale network of discrete interacting regions has proved fruitful in the analysis and modelling of neural dynamics [1]. Moreover, this abstraction lends neuroscientists the language and tools of statistical physics in the hope of uncovering the central mechanisms driving brain function and their links to observed neural dynamics [2, 3]. For instance, recent data captured by functional imaging showed large scale violations of detailed balance in human brain dynamics, suggesting that the brain is operating far from equilibrium [4]. This fundamental observation has prompted the development of a range of techniques to provide a measure for the degree of non-equilibrium in neuroimaging time-series recorded in different conditions [5–10]. These measures have shown that the degree of non-equilibrium is elevated during cognitive tasks [4–7] whilst reduced in both impairments of consciousness [11], sleep [10] and Alzheimer’s disease [12], indicating that non-equilibrium may be a key signature of healthy consciousness and cognition in the brain [13]. Despite this, current methods are restricted to aggregate measures of non-equilibrium. We present a novel approach to the analysis of non-equilibrium brain dynamics that is able to measure the irreversibility of individual, higher-order interactions to gain valuable insight into the organisation of neural dynamics.

The second law of thermodynamics asserts that, in the absence of entropy sinks, the average entropy of a system increases as time flows forwards [14, 15]. More specifically, a system at a steady-state dissipating heat to its environment causes an increase in entropy [16, 17]. This results in the system breaking the detailed balance condition and results in an asymmetry in the probability of transitioning between system states [18]. This, in turn, yields macroscopically irreversible trajectories from reversible microscopic forces inducing what Eddington denoted ‘the arrow of time’ (AoT) [19]. The rate at which a system dissipates entropy, the ‘entropy production rate’ (EPR), is a natural measure of the degree of non-equilibrium in the stationary state, as it is zero in equilibrium and positive out of equilibrium [20]. Results in modern non-equilibrium thermodynamics have shown that the EPR of a non-equilibrium system can be derived from the irreversibility of observed trajectories [21–25]. In particular, the EPR is given by,

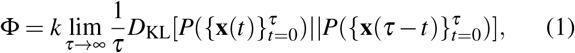

where 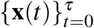 and 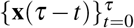 represent a trajectory and its time-reversal, *P*(*·*) represents the ‘path probability’, the probability of observing that specific trajectory, *k* is Boltzmann’s constant, and *D*_KL_ represents the Kullback-Leibler divergence (KLD),

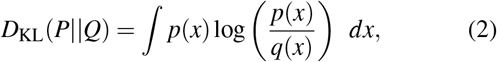

which measures the distance between two probability distributions *P* and *Q* with densities *p* and *q* respectively [24, 25]. In the case of real-world data, trajectories are sampled at discrete time-points forming a multivariate time-series (MVTS), and the EPR is lower-bounded by the irreversibility of the observed MVTS. As a result, the irreversibility of a neural recording is a natural measure of the degree to which the neural dynamics are out of equilibrium [13].

Two complimentary interpretations of the AoT in the brain have been given. First, the hierarchical organisation of positions in state-space, that results from asymmetrical transition probabilities, has been linked to the dynamic hierarchical organisation of brain regions [7, 26, 27]. Second, the AoT has been interpreted as inducing a ‘causal flow’ in the system where some regions emerge as information ‘sources’ and others as ‘sinks’ with these relationships identifiable from irreversibility analysis [7, 8]. These studies for quantifying non-equilibrium in the brain approximate the global evidence for the AoT in time-series using techniques such as estimating transitions between coarse-grained states [4], with time-shifted correlations [5], machine learning [6] or with model-based approaches [7–10]. However, the AoT and the corresponding production of entropy is a macroscopic property of the system, emerging from interactions between the microscopic variables at multiple scales. Recent theoretical research has shown that the AoT can be decomposed into unique contributions arising at each scale within the system [28, 29] or into spatiotemporal modes of oscillation [30], offering insights beyond a global level of non-equilibrium in the brain. Motivated by these insights, we present the Directed Multiplex Visibility Graph Irreversibility (DiMViGI) framework, as illustrated in Fig. 1, for analysing the irreversibility of multivariate signals at multiple levels using network analysis of time-series, in particular the visibility graph [31, 32]. Using the DiMViGI framework, we investigate the irreversibility of human brain signals, captured by magnetoencephalography (MEG), during a long-term recognition task of musical sequences that utilised long-term memory [33–39]. Our analysis covers all possible levels in the system and is able to capture the higher-order organisation of brain regional interactions yielding interpretable and novel insights into the neural dynamics underpinning long-term memory and auditory recognition.

**FIG. 1.**
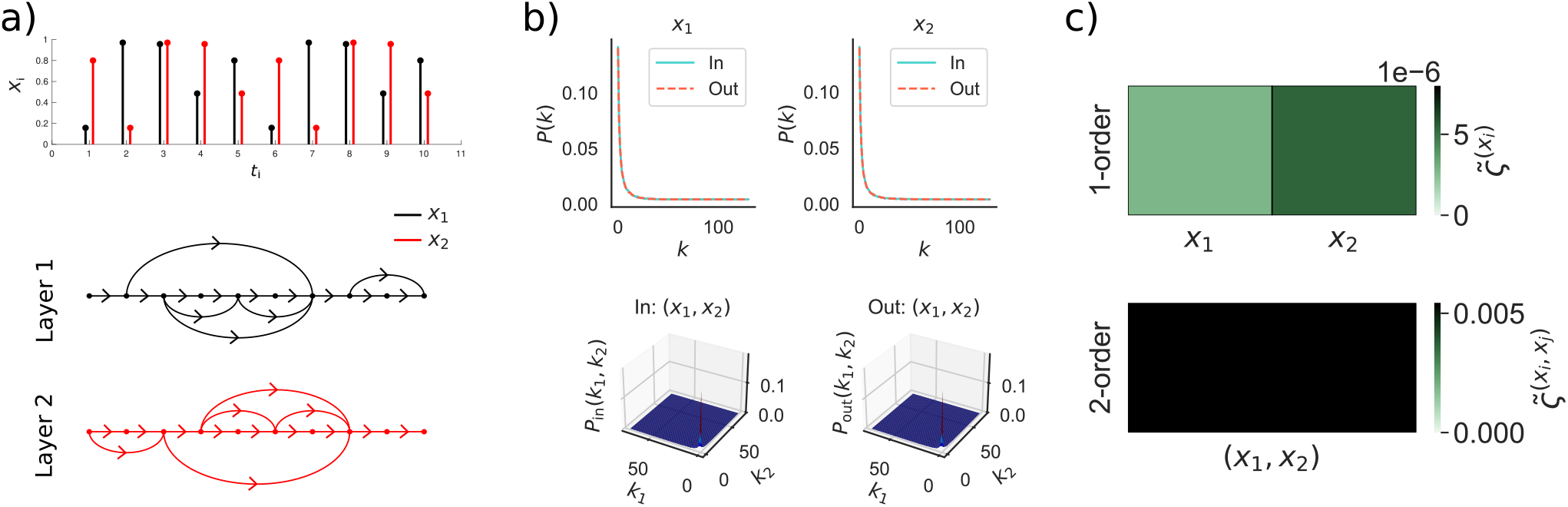
The DiMViGI workflow. The method is able to measure the irreversibility of each interaction in a multivariate time-series. It is comprised of three stages, illustrated here with a random time-series of 2 variables: (a) First, we construct a 2-layer directed multiplex visibility graph from the multivariate time-series where each layer represents a variable and each node represents a time-point. The connections are made according to the visibility criterion defined in Eq. 7 and illustrated in Fig. 2. (b) Second, we calculate the in- and out-degree distributions for each tuple at each level. In the 2-variable system, there are 3 such tuples: the singletons, (*x*_1_), (*x*_2_) and the pair (*x*_1_, *x*_2_). The top left/right panels show the in- and out-degree distributions for the singletons (*x*_1_), (*x*_2_) respectively. The bottom two panels show the in- (Left) and out- (Right) degree distribution of the pair (*x*_1_, *x*_2_). (c) Third, we measure the Jensen-Shannon divergence of the in- and out-degree distributions for each tuple in the system. We show the 1-order irreversibility, 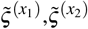, of the singletons (*x*_1_), (*x*_2_) (top) and the 2-order irreversibility, 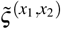, of the pair (*x*_1_, *x*_2_) (bottom).

### QUANTIFYING THE ARROW OF TIME IN MULTIVARIATE INTERACTIONS

As the evidence for the AoT can be inferred from the irreversibility of observed trajectories, we focus on the quantity,

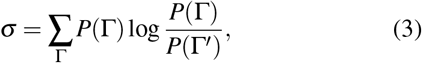

where Γ is a stochastic trajectory, Γ*′* is its time-reversal and *P*(Γ) is the probability of observing that specific trajectory. Eq. 3 is precisely the KLD between the forward and back ward path probabilities, which is a natural measure of the irreversibility of a stochastic process [23]. Inspired by previous decompositions [28, 29], we note that individual interactions can have differential levels of irreversibility within a globally non-equilibrium system. Our framework aims to compute the irreversibility of individual *k*-tuples of variables in a MVTS in order to compare interactions at each level, defined by *k*. Firstly, we consider the projection of an *N*−dimensional trajectory, 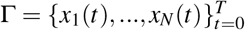, into the portion of state-space defined by the *k*-tuple of variables 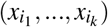, to be the *k*-dimensional trajectory,

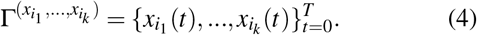

The DiMViGI framework then quantifies the marginal irreversibility of a given tuple by approximating,

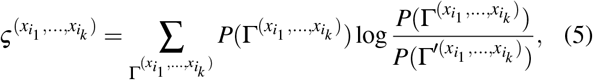

using visibility graphs, as will be detailed subsequently. As a result, we are able to identify tuples of variables whose multivariate trajectory is highly irreversible indicating a strongly non-equilibrium interaction between the variables in this tuple, which also suggests the presence of a hierarchical structure within the tuple [7].

### MEASURING IRREVERSIBILITY WITH THE MULTIPLEX VISIBILITY GRAPH

We build on the growing paradigm of network analysis of time-series that has gained traction in the analysis of neural signals [40, 41]. These methods are characterised by mapping a time-series into a corresponding network. For instance, the visibility algorithm maps a univariate time-series into a so-called ‘visibility graph’ (VG) [31]. VGs and their variations are a powerful model-free tool for mapping a continuous-valued time-series into a discrete object. Their versatility, as well as their lack of assumptions on the underlying dynamics, has lent them to diverse applications, in particular in neuroscience [40, 41], as well as in the calculation of information-theoretic quantities from complex and chaotic dynamics [42]. Explicitly, given a time-series {*X*_*i*_}_*i*∈*I*_ with time indices {*t*_*i*_}_*i*∈*I*_, where *X*_*i*_ ∈ ℝ and *I* is the index set, the VG has one node for each *i* ∈ *I*. Nodes *i, j* ∈ *I* are connected by an edge if the corresponding data-points (*t*_*i*_, *X*_*i*_) and (*t* _*j*_, *X*_*j*_) are ‘mutually visible’ i.e. that they satisfy that, for any intermediate data-point (*t*_*k*_, *X*_*k*_) with *t*_*i*_ *< t*_*k*_ *< t* _*j*_,

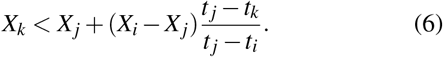

In geometric terms, this condition is met if (*t*_*i*_, *X*_*i*_) is visible from (*t* _*j*_, *X*_*j*_). That is, the line connecting (*t*_*i*_, *X*_*i*_) and (*t*_*j*_, *X*_*j*_) does not cross any intermediate data-points as shown in Panel b) of Fig. 2. Trivially, each node is connected to its neighbours whilst large positive fluctuations become hubs with many connections due to their greater visibility. This construction can be naturally extended to a MVTS by considering the ‘multiplex visibility graph’ (MVG) [43]. Given a MVTS with *N* variables, the MVG is a multi-layer graph, a so-called ‘multiplex’, with *N* independent layers with the same node base. Applying the visibility algorithm to each variable in turn yields a series of VGs which each define one layer of the MVG.

**FIG. 2.**
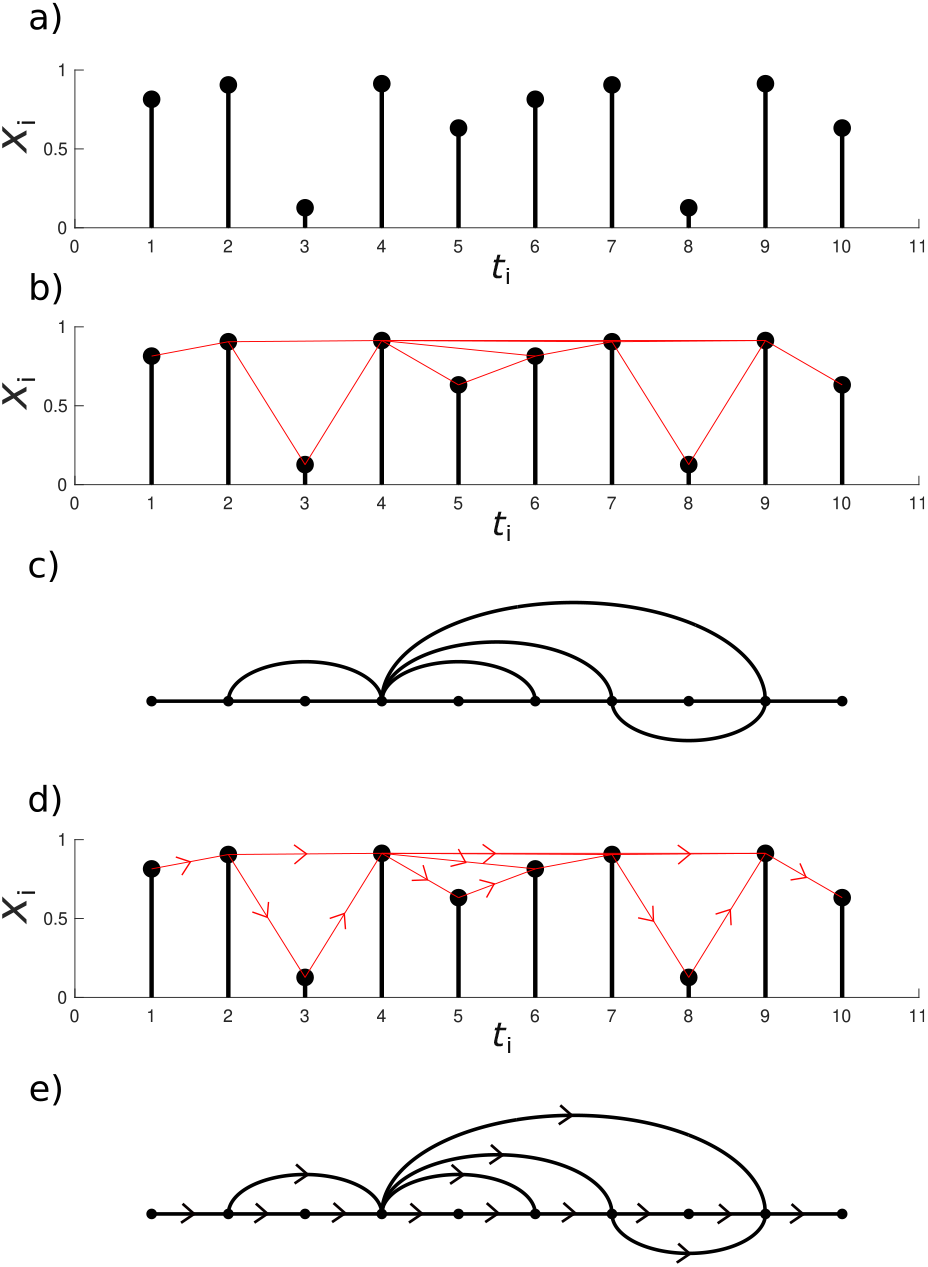
Visibility graphs. An example of a visibility and a directed visibility graph constructed from a random time-series. (a) A random equi-spaced time-series. (b) The red lines connected data points that mutually visible. (c) The visibility graph associated with the random series. (d) A time-series showing visibility directed forward in time. (e) The directed visibility graph corresponding to the above series.

We can further generalise the VG to measure irreversibility in univariate time-series by extending the undirected VG to a time-directed counterpart (DVG) [32, 44]. To do so, we simply direct the edges ‘forward in time’. For example, an edge connecting time-points *t*_*i*_ *< t* _*j*_ is now directed *i* → *j* (see Panels d-e) of Fig. 2). We then decompose the degree *d* of a node into the sum of the in-going and out-going degree,

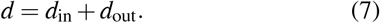

A univariate stationary process, *X* (*t*), is time-reversible if the trajectory {*X* (*t*_1_), …, *X* (*t*_*T*_)} is as probable as {*X* (*t*_*T*_), …, *X* (*t*_1_)} [45]. Therefore, in the case of a reversible process, the in- and out-going degree distributions of the associated DVG should converge [32, 44]. It follows that the level of irreversibility can be captured by measuring the divergence between the in- and out-going degree distributions. We extend this method to the case of MVTS. We direct the edges of the MVG such that they go forward in time yielding a directed MVG (DMVG). Since this is a multiplex graph, we can calculate the multivariate joint, over all layers, in- and out-going degree distributions, and all associated marginals.

Explicitly, we consider a MVTS with *N* variables and *T* time points, given by {**X**(*t*_1_), …, **X**(*t*_*T*_)}, where **X**(*t*_*i*_) = (*x*_1_(*t*_*i*_), …, *x*_*N*_ (*t*_*i*_)) ∈ ℝ^*N*^ and construct its associated DMVG. For a given *k*-tuple of variables, (*n*_1_, …, *n*_*k*_), we calculate the multivariate marginal in-going and out-going degree distributions:

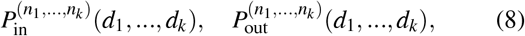

where 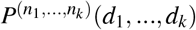 is the probability of a node having degree *d*_*i*_ in layer *n*_*i*_ for all *i* simultaneously. We then compute the divergence between these particular in- and out-going marginal distributions using Jensen-Shannon divergence (JSD) (see Materials and Methods) to obtain a measure of the *k*-order irreversibility,

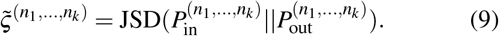

As we are considering the multivariate joint distribution, we are quantifying irreversibility in the multivariate state-space. Repeating this for all possible *k*-tuples in the system, we quantify the relative irreversibility of each interaction at a given level. We can repeat this process for all values of *k*, thus measuring irreversibility at all levels.

In summary, the DiMViGI framework, shown in Fig. 1, begins with a MVTS of neural activity. The series is mapped into the associated DMVG using the visibility algorithm. We calculate the joint in and out-degree distributions and all the possible marginal in- and out-degree distributions. We measure the JSD between the pairs of in- and out-marginals for each tuple in the system to quantify the irreversibility of that interaction. At each level *k*, we can then compare the relative irreversibility of each *k*-order interaction to identify the dominant irreversible interactions.

### ANALYSIS OF MEG DURING LONG-TERM RECOGNITION

We consider MEG recordings from 51 participants with 15 trials per participant source-localised into 6 regions of interest (ROIs) collected according to the experimental paradigm presented in Fig. 3, described in Materials and Methods, SI and in Ref. [33]. The ROIs include the auditory cortices in the left and right hemispheres (ACL, ACR); the hippocampal and inferior temporal cortices in the the left and right hemispheres (HITL, HITR) and two medial regions, the bilateral medial cingulate gyrus (MC) and the bilateral ventro-medial prefrontal cortex (VMPFC). Panel a) of Fig. 4 shows a schematic representation of the regions. The participants performed an auditory recognition task during the MEG recordings (Panel a), Fig. 3). First, they memorised a short musical piece. Next, they were presented musical sequences and were requested to state whether the sequence belonged to the original music or was a varied version of the original sequences. Since differences between experimental conditions have been described in detail by Bonetti et al [33] and are beyond the scope of this work, here, we consider only one experimental condition, where participants recognised the original, previously memorised sequences.

**FIG. 3.**
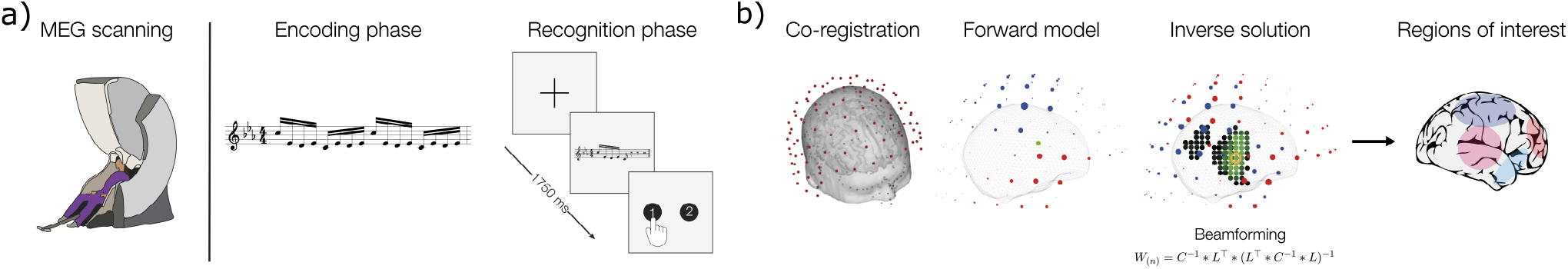
Experimental paradigm for the collection and processing of MEG data. (a) The brain activity in 51 participants was collected using magnetoencephalography (MEG) while they performed a long-term auditory recognition task. Participants memorised a 5 tone musical sequence. They were then played 5 further sequences of tones that were either the original sequence or a modified version. They then were requested to state whether the sequence belonged to the original music or was a varied version of the original sequences. In this analysis we only consider the experimental condition where participants were played the original memorised sequence. (b) The MEG data was co-registered with the individual anatomical MRI data, and source reconstructed using a beamforming algorithm. This procedure returned one time-series for each of the 3559 reconstructed brain sources. Six main functional brain regions (ROIs) were derived. The neural activity for each ROI was extracted yielding a multivariate time-series. For further details on the experimental set-up see Materials and Methods and SI. For a comparison between experimental conditions see Bonetti et al [33].

**FIG. 4.**
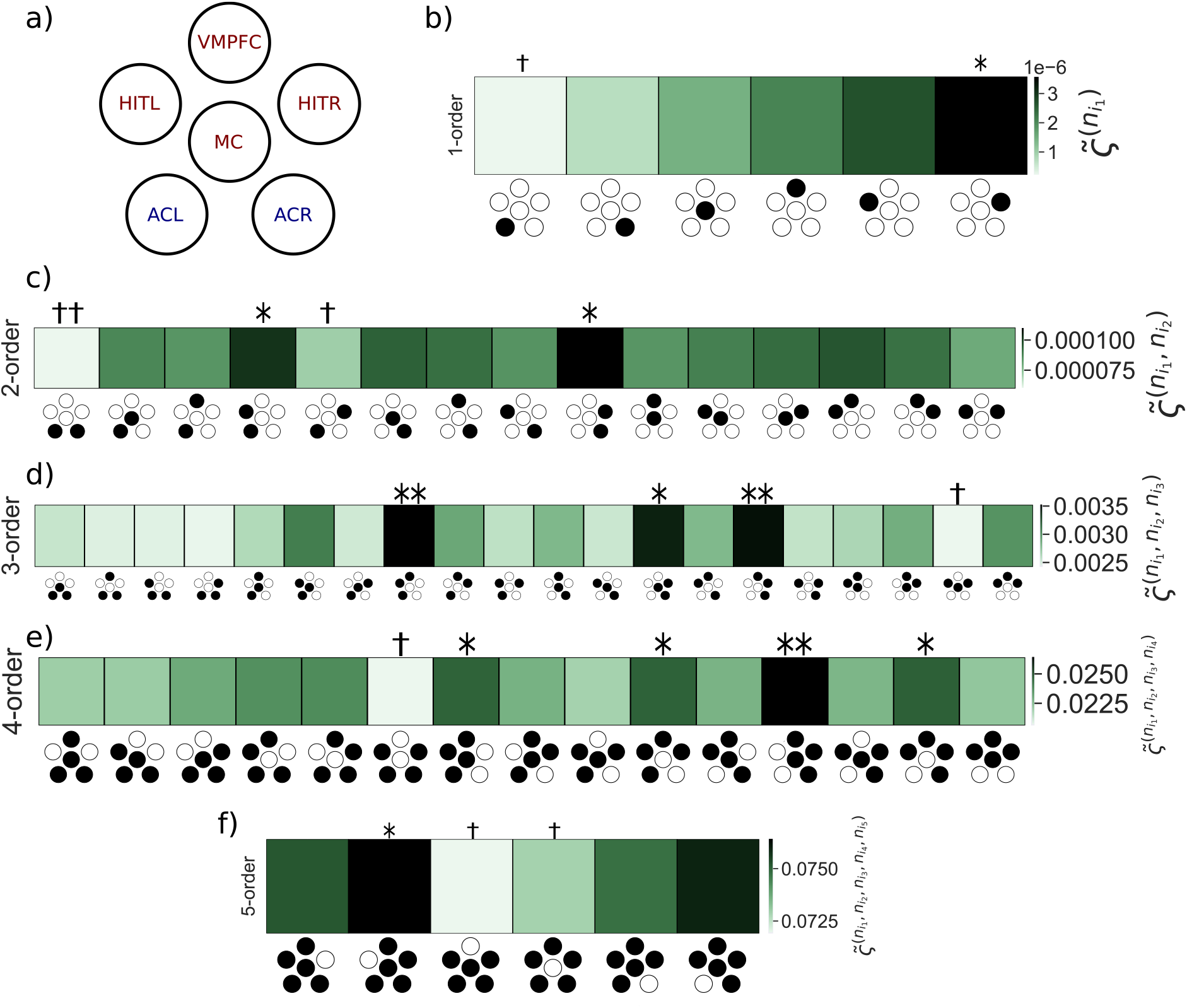
DiMViGI analysis of 6-ROI MEG recordings during a long-term memory task. The number of (*)/(†) represents the number of standard deviations above/below the mean for a particular tuple at that level. (a) Schematic diagram showing the organisation of the ROIs in the MEG recordings. The ROIs are ACL/R: auditory cortex left/right; MC: medial cingulate gyrus; VMPFC: ventro-medial prefrontal cortex; HITL/R: hippocampal inferior temporal cortex left/right. Cognitive regions are in red and sensory regions in blue. (b) 1-order irreversibility at cohort-level. At this level, we consider irreversibility of each signal in isolation. The hippocampal regions are the most irreversible whilst the sensory regions are the most reversible. (c) 2-order irreversibility at cohort-level. The pairs that show the most irreversibility are those that include a sensory and hippocampal pair in the same hemisphere (ACL/R, HITL/R). The most reversible pair is (ACL, ACR) which is made up of two sensory regions. (d) 3-order irreversibility at cohort-level. The triplets that are most irreversible are those that include an intra-hemispheric sensory and hippocampal pair as well as the prefrontal cortex (ACL/R, HITL/R, VMPFC). The most reversible contains both hippocampal regions and the medial cingulate gyrus, (HITL, HITR, MC). (e) 4-order irreversibility at cohort-level. The quadruplets that are most irreversible are those that include a hippocampal and sensory pair and both medial regions (ACL/R, HITL/R, MC, VMPFC) and those that include both hippocampal regions, a sensory region and the VMPFC. The most reversible is the quadruplet that contains no medial regions. 5-order irreversibility at cohort-level. The most reversible quintuplets are those that omit a medial region, in particular the quintuplet that omits the VMPFC.

For each participant and trial, we construct the DMVG. Next we estimate every marginal in- and out-degree distribution using each DMVG as a sample and calculate the JSD. We denote the JSD between *k*-dimensional degree distributions as the *k*-order irreversibility. Alternatively, for each participant in isolation, the degree distributions can be calculated using only their associated trials to get an estimate of the *k*−order irreversibility for each participant and each tuple (see SI). However, due to the higher number of samples, the cohort-level analysis is more robust and hence is our focus in this report. The results of the DiMViGI analysis are presented in Figure 4. We note that the darker colours represent tuples with greater irreversibility whilst the lighter colours reflect more reversible interactions. The icon along the *x*-axis indicates which tuple is being considered, with reference to the schematic in Panel a) of Fig. 4, with the included regions coloured in black. Furthermore, we highlight statistically significant tuples at each level. The number of (*)/(†) indicates the number of standard deviations above/below the *k*−level mean.

We begin our analysis at 1-order. Whilst individual (microscopic) variables are often reversible in a non-equilibrium complex system, the ROIs considered here reflect a very coarse parcellation of the brain. At this level, we are considering each ROI, which is composed of many truly microscopic variables, in isolation and note that each one shows significant irreversibility. It is clear from Panel b) of Fig. 4, that the ROIs have a clear disparity in their levels of irreversibility. The sensory ROIs are more reversible than the medial and hippocampal ROIs. Furthermore, there is a skew towards the right hemisphere being more irreversible than the left. This result emerges consistently across all levels. Next, we consider the irreversibility of pairwise interactions (*k* = 2). Panel c) of Fig. 4 shows the 2-order irreversibility for all pairs. We are able to identify strongly irreversible pairs such as the intra-hemispheric pairs (ACL, HITL) and (ACR, HITR). On the other hand, cross-hemispheric pairs, e.g. (ACL, ACR), are the most reversible, indicating a lack of interaction between them. The strong hemispheric symmetry in the results validates the findings, as it is an expected and intuitive observation. Panel d) of Fig. 4 shows the irreversibility for each triplet interaction in the system. The highly irreversible triplets are those that include a hemispheric pair alongside a medial region, with those containing the VMPFC, a region known to drive brain dynamics during task [46], being particularly irreversible. Panel e) of Fig. 4 shows that the most irreversible quadruplet interactions are composed of a hemispheric pair alongside both medial regions as well as those that contain (VMPFC, HITL, HITR) alongside a sensory region. Conversely, the quadruplet containing no medial regions, is the most reversible, and therefore has the least interaction. This is particularly interesting as this quadruplet is made up of the two most irreversible pairs yet they do not appear to interact as a foursome. Therefore, this framework is truly capturing higher-order interactions that cannot simply be decomposed into a sum of independent interactions of lower order. Finally, Panel f) of Fig. 4 shows that quintuplets that contain both medial ROIs are the most irreversible. Furthermore, the quintuplet that does not contain the VMPFC has the most reversible interaction. Whilst we have attempted to interpret the results from the perspective of the hierarchical and higher-order organisation of the auditory system, we note that outliers would be expected to arise naturally due to statistical variation. Nevertheless, due to the consistency of our results across levels, for example the hemispheric symmetry that is observed at each level, such results cannot be explained purely by chance. Furthemore, a sub-sampling analysis shows that the error in irreversibility measurements are typically smaller than differences between tuples implying a range of statistically significant differences (see SI).

We can interpret this result in the context of predictive coding and its links to sensory tasks [47–49], as well as through the hierarchical organisation of the auditory system. The participants are exposed to a memorised tonal sequence that does not deviate from their expectation of what they were about to hear. Under the theory of predictive coding, this would result in an adjustment of a participant’s prior expectations, facilitated by asymmetric, hierarchical interactions between brain regions at multiple levels, in order to reinforce the prior expectations in light of the new sensory information [50]. This in turn would lead to a cascade of interactions between key ensembles of regions whose function is optimised for the process of auditory recognition. As irreversible brain dynamics stem from irreciprocal and hierarchical interactions, such a mechanism results in marked irreversibility in the emergent dynamics [7].

## DISCUSSION

In this study, we describe a novel framework for measuring the emergence of non-equilibrium dynamics, through multivariate irreversibility, at multiple system levels. We are able to capture the irreversibility of each possible interaction in a MVTS of signals. Applying the DiMViGI framework to neural recordings obtained during a long-term memory recognition task, we investigate the higher-order organisation, and the associated non-equilibrium interactions, of brain regions and how they break time-reversal symmetry during an auditory recognition task. The results clearly show a broad distribution of irreversibility at each system level; hence we are able identify which interactions are particularly irreversible, which we interpret as a correlate of a hierarchical and synergistic interaction. Furthermore, we link irreversibility to hierarchical predictive coding and theorise that non-equilibrium interactions could emerge as a consequence of the modulation of prior expectations in light of new sensory information [50]. According to the theory of predictive coding, this might be realised through hierarchically asymmetric interactions that, in turn, induce the emergence of irreversibility at multiple system levels [7, 51, 52]. Within this context, the DiMViGI framework confirms the hierarchical organisation of the auditory system [53–56], with reciprocal connections, such as those found within the auditory cortex, resulting in more reversible dynamics, and hierarchical relationships, such as those found between the auditory cortex and the hippocampus, resulting in markedly irreversible dynamics. Furthermore, our approach goes beyond typical approaches to the auditory system, such as the analysis of co-activation and functional connectivity [57, 58] or the identification of cortical-gradient hierarchies [33, 55], by uncovering higher-order interactions within the auditory system between triplets and quadruplets of brain regions. In particular, at higher-orders, irreversibility reveals synergistic interactions between hippocampal, cingulate gyrus and sensory regions for the distributed processing required for audition and long-term recognition. As a result, our approach yields insights that offer a new perspective on the flow of information during audition. Whilst a recent analysis of these neural recordings with standard methods was able to identify a hierarchy of information processing in the brain during long-term recognition [33], the introduction of the DiMViGI framework appears crucial to uncovering the higher-order and non-equilibrium nature of the interactions. Such insights are opaque to traditional analyses but emerge from the unique lens of non-equilibrium statistical physics.

The implications of the framework and the associated results are multi-fold. Firstly, we go beyond aggregate [4–7, 9, 10] or univariate [32, 44] measures of irreversibility, expanding the exisiting quiver of techniques for studying non-equilibrium in the brain to include a multilevel approach.

Our technique is able to capture differences in irreversibility across scales in continuous time-series, inspired by recent theoretical work for binary variables [28, 29], that is non-specific and can be applied to MVTS from any domain to identify particular highly non-equilibrium interactions. Our approach differs from Refs. [28, 29] as we do not attempt to measure the unique contribution to the AoT of a specific *k*-body interaction by discounting the irreversibility of all sub-interactions contained within the tuple. Instead, we measure the irreversibility of the tuple as a whole. In Section 6 of the SI we consider an extension of our approach to relate our framework more closely to the approach of Refs. [28, 29], by measuring the unique contribution of each *k*−body interaction, defined recursively as,

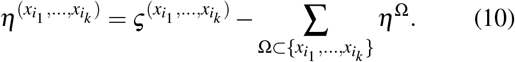

However, we note that the exact decomposition of the EPR presented in Refs. [28, 29] relates to discrete, Markovian and multi-partite dynamics and thus does not apply directly to continuous MVTS. Moreover, in Section 5 we show that irreversibility in our method only decomposes in the case of independent variables.

Our framework builds on the sustained interest in identifying higher-order interactions in neural recordings and other MVTS [59–63], particularly in information theoretic analyses of brain data that reveal how higher-order functional interactions shape neural dynamics [64–66]. Notably, many higher-order frameworks are either computationally, or by formulation, restricted to studying either triplet [60, 61, 63, 64] or system-wide interactions [59], whilst our results extend easily to all possible levels in the system. Our framework attempts to bridge the broader discussion on higher-order mechanisms and behaviours in complex systems [67–69] with techniques from non-equilibrium thermodynamics [20] through the quantification and interpretation of multilevel irreversibility. Finally, our work further solidifies the visibility algorithm, and network analysis of time-series, as an empirically useful tool in the analysis of neural data [40, 70].

Despite these promising results, we note some nuanced limitations in our framework. Whilst the visibility algorithm and the degree distribution approach reduces the dimension of the data, we are still computing an entropy between high-dimensional distributions which is computationally restrictive. This can be circumvented limiting the support of the degree-distribution to exponentially improve computational efficiency whilst minimally affecting numerical accuracy (see SI). Nevertheless, analysing all possible interactions yields a combinatorial explosion, hence we opt for a coarse, low-dimensional, parcellation of the brain that allows us to analyse the system at all possible levels. However, the highlighting of individual tuples is most meaningful when there is a strong intuition about the nature of the interaction, which can be only be expected in low-dimensional parcellations where ROIs are clear, functionally-segregated brain areas.

Additionally, we note that our measure is undirected within the tuple, meaning we cannot identify the direction of information flow as one can with classical measures of causality [71, 72] or some approaches to the AoT [7, 8]. However, we note that the AoT represents directed flow between states and not variables, meaning it is not a direct measure of causality, but instead capturing a distinct, but related, phenomena in interacting dynamics. Finally, measuring the irreversibility of finite-length time-series naturally induces a bias due to the finite sampling of the state-space [4, 29]. In order to validate that the measured irreversibility emerges from non-equilibrium dynamics and not from finite-data errors, we employed both surrogate-testing using shuffled time-series and sub-sampling approaches to validate the significance of our results (see SI Section 4).

A key advantage of the DiMViGI framework is the ability to scale between levels with a consistent approach. Strictly local measures such as auto- and cross-correlations are limited to individual and pairwise interactions [73, 74]. On the other hand, simply applying global measures to each subset of variables in the time-series, such as coarse-graining or using a model-based measure, yields an inconsistent approach where different tuples cannot be compared fairly. Our framework extends consistently to all levels thus yielding directly comparable quantities at each level.

## CONCLUSIONS

In this work, we have introduced the Directed Multiplex Visibility Graph Irreversibility framework for measuring the irreversibility of multivariate interactions at all levels within a system. We applied this method to neural recordings during a long-term auditory recognition task to study the relative irreversibility of different interactions between brain regions. Doing so, we were able to demonstrate the hierarchical, higher-order organisation of brain dynamics during tasks. This analysis suggests that reinforcement of prior expectations during an auditory recognition task is facilitated through a hierarchy of irreversible higher-order interactions in the brain, an observation that we link to both the mechanisms of predictive coding and the hierarchical structure of the auditory system. Furthermore, we highlighted the particular combinations of cognitive and sensorial regions that are preferentially recruited during audition and long-term recognition. This framework is non-specific and provides a general tool for investigating higher-order interactions and non-equilibrium dynamics in MVTS emerging from other complex systems.

## MATERIALS AND METHODS

### Estimating degree distributions from finite samples

For each sample, a MVTS, we construct the DMVG, defined by the multiplex adjacency matrix, *A*,

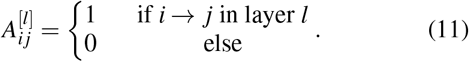

Then we calculate the in- and out-degree of each node in each layer

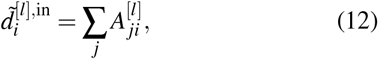

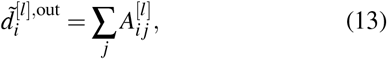

where 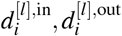 are the in-and out-degree of node *i* in layer *l* respectively.

For a *k*−tuple (*n*_1_, …, *n*_*k*_), we calculate 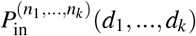 by counting the number of nodes *i*, across all samples, where

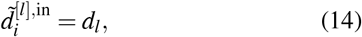

for each *l* ∈ {1,…, *k*} simultaneously and for *d*_*l*_ ∈ {1,…, *d*_max_} where *d*_max_ is the maximum degree of a node in the multi-layer graph, and then dividing through by the total number of nodes in all samples. We calculate the same for 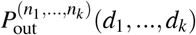.

As we are using a finite number of samples, we then perform Laplace smoothing to eliminate singularities of the form *P*(*x*) = 0 *< Q*(*x*) for which divergence is ill-defined. Instead of using,

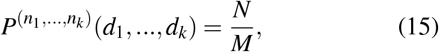

where *N* is the number of nodes satisfying condition 14 and *M* is the total number of nodes across samples, we perform the following replacement,

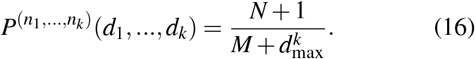

Such an approach is equivalent to assuming a uniform Bayesian prior for the degree distributions [75].

### Computing Jensen-Shannon divergence

We quantify the divergence between the in- and out-degree distributions using Jensen-Shannon divergence (JSD) which is a symmetrised version of Kullback-Leibler divergence (KLD) that does not suppose a model-data relationship [76]. This is defined between two probability distributions *P, Q* as

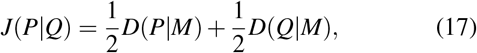

where 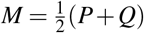 is an averaged distribution and *D*(*·*) represents the KLD, given by,

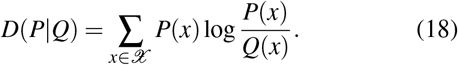

As 𝒳 represents the support of the distribution, it takes the form {1,…, *d*_max_}^*k*^ where *k* is the dimension of the probability distributions and *d*_max_ is the maximum degree of a node in the multi-layer graph. For computational feasibility, *d*_max_ can be limited during the calculation of JSD, truncating the sum. For 5-order analysis, we limit *d*_max_ to 75. For a systematic analysis of the effect of degree limiting see SI.

### Magnetoencephalography (MEG) data

#### Participants

The participant cohort consisted of 83 healthy volunteers made up of 33 males and 50 females with ages in the range 18 to 63 and a mean age of 28.76 *±* 8.06. The 51 participants included in this analysis included 22 males and 29 females with ages in the range 18 to 63 and a mean age of 27.57 *±* 7.13. Participants were recruited in Denmark, came from Western countries, reported normal hearing and gave informed consent before the experiment. The project was approved by the Institutional Review Board (IRB) of Aarhus University (case number: DNC-IRB-2020-006) and experimental procedures complied with the Declaration of Helsinki – Ethical Principles for Medical Research. After pre-processing, the 51 participants with at least 15 non-discarded trials in the first experimental condition were included in the analysis. Only trials where participants correctly identified the sequence were included. For those participants with more than 15 trials, 15 trials were randomly sampled.

#### Experimental stimuli and design

We employed an old/new paradigm auditory recognition task [33, 35, 36, 38]. Participants listened to a short musical piece twice and asked to memorise it to the best of their ability. The piece was the first four bars of the right-hand part of Johann Sebastian Bach’s Prelude No. 2 in C Minor, BWV 847. Next, participants listened to 135 five-tone musical sequences, corresponding to 27 trials in 5 experimental conditions, of 1750 ms each and were requested to indicate if the sequence belonged to the original music or was a variation. Differences between experimental conditions have been described in detail by Bonetti et al [33]. We consider one experimental condition, where participants recognised the original, previously memorised sequences.

#### Data acquisition

MEG recordings were taken in a magnetically shielded room at Aarhus University Hospital, Aarhus, Denmark using an Elekta Neuromag TRIUX MEG scanner with 306 channels (Elekta Neuromag, Helsinki, Finland). The sampling rate was 1000 Hz with analogue filtering of 0.1-330 Hz. For further details on the data acquisition see SI.

#### MEG pre-processing

First, raw MEG sensor data was processed by MaxFilter [77] to attenuate external interferences. We then applied signal space separation (for parameters see SI). Then the data was converted into Statistical Parametric Mapping (SPM) format, preprocessed and analyzed in MATLAB (MathWorks, Natick, MA, USA) using in-house codes and the Oxford Centre for Human Brain Activity (OHBA) Software Library (OSL) [78]. The continuous MEG data was visually inspected and large artefacts were removed using OSL. Less than 0.1% of the collected data was removed. Next, independent component analysis (ICA) was implemented to discard artefacts in the brain data from heart-beats and eye-blinks (for details see SI) [79]. Lastly, the signal was epoched in 135 trials, 27 trials for each of 5 experimental conditions and the mean signal recorded in the baseline (the post-stimulus brain signal) was removed. Each resulting trial lasted 4400 ms plus 100 ms of baseline time.

#### Source reconstruction

We employed the beamforming method to spatially localise the MEG signal [80]. For details on the beamforming algorithm and the implementation see SI.

## Code and data availability

The code used to implement the DiMViGI framework will be made available at https://github.com/rnartallo/multilevelirreversibility upon acceptance.

The in-house code used for MEG pre-processing is available at https://github.com/leonardob92/LBPD-1.0.

The multimodal neuroimaging data analysed here is available upon reasonable request.

## AUTHOR CONTRIBUTIONS

R.N.K designed and performed the analysis and wrote the manuscript. L.B. designed the analysis and the experiment, collected and pre-processed data, and edited the manuscript. G.F.R. designed the experiment and collected and preprocessed the data. P.V. designed and secured funding for the experiment. G.D. edited the manuscript. M.L.K, R.L. and A.G. supervised the research, analysed the data and edited the manuscript.The authors declare no competing interests.

## ACKNOWLEDGMENTS

R.N.K was supported by an EPSRC doctoral scholarship from grants EP/T517811/1 and EP/R513295/1. L.B, M.L.K and P.V were supported by the Center for Music in the Brain (MIB), funded by the Danish National Research Foundation (project number DNRF117). L.B. was also supported by Carlsberg Foundation (CF20-0239), Lundbeck Foundation (Talent Prize 2022), Linacre College of the University of Oxford (Lucy Halsall fund) and Nordic Mensa Fund. G.F.R. was supported by Fundación Mutua Madrileña. G.D. was supported by the AGAUR research support grant (ref. 2021 SGR 00917) funded by the Department of Research and Universities of the Generalitat of Catalunya, by the project NEurological MEchanismS of Injury, and the project Sleep-like cellular dynamics (NEMESIS) (ref. 101071900) funded by the EU ERC Synergy Horizon Europe and by the project PID2022-136216NB-I00 financed by the MCIN /AEI /10.13039/501100011033 / FEDER, UE., the Ministry of Science and Innovation, the State Research Agency and the European Regional Development Fund. M.L.K was further supported by Centre for Eudaimonia and Human Flourishing funded by the Pettit and Carlsberg Foundations. R.L. was supported by the EPSRC grants EP/V013068/1 and EP/V03474X/1.

## Supplementary Information

### I. INTRODUCTION

In this supporting information, we provide additional results and analysis not included in the main manuscript. This supporting information is organised as follows. In Section II, we show the results of the directed multiplex visibility graph irreversibility (DiMViGI) framework applied to data from individual participants to produce distributions of participant spread for the irreversibility of each tuple, rather than the cohort-level analysis presented in the main manuscript. We assess the significance of the differences between tuples using pairwise *t*−tests and one-way ANOVAs. Moreover, we calculate the correlations between the cohort and participant level analysis. In Section III, we further validate the significance of the results obtained at the participant level by shuffling the time-series to produce surrogate data. We show that shuffling restores detailed balance and that the irreversibility of the true signals is significantly higher than the surrogate data. This indicates that the measured irreversibility is due to the non-equilibrium dynamics present in the time-series and not bias from finite data. In Section IV, we employ a second method to correct for biases stemming from ‘finite’ data. We use sub-sampling to estimate the irreversibility for various fractions of our data. We show that our results are robust to various sub-samplings of the data. Additionally, this produces a distribution of the irreversibility measurement, which we use to validate that the relative error in our measurement is significantly lower than the difference between tuples. Next, in Section V, we show that the DiMViGI framework factorises for independent variables, theoretically validating the argument that it captures ‘true’ higher-order interactions. Using the factorisation, in Section VI, we are able to define the unique irreversibility generated by a higher order interaction by removing the lower-level interactions. We compare this to the ‘combined’ results presented in the manuscript. In Section VII, we validate that our method captures a correlate of the entropy production rate by using simulated data from four specific examples of the multivariate Ornstein-Uhlenbeck process. In Section VIII, we investigate the effect on the results of limiting the maximum degree in the support of the distributions, an approach that improves computational efficiency whilst only minimally reducing accuracy. In Section IX, we discuss the definitions of entropy production rate for Markovian and non-Markovian dynamics. In Section X we present a comprehensive description of the experimental paradigm and the techniques used to record and pre-process the magnetoencephalography (MEG) data. In Section XI, we present an evaluation of the signal quality and signal to noise ratio as well as visualisations of baseline activity. Finally, in Section XII, we compare the results of applying our analysis to the full epoch, as presented in the paper and the rest of the SI, as well as an epoch with the pre-stimulus baseline removed and an epoch with both the pre-stimulus and post-task reset to baseline removed. We use quadruplets as an example, and show that the results are equivalent.

### II. THE DIMVIGI FRAMEWORK APPLIED TO PARTICIPANT-LEVEL DATA

In this section we show the results of applying the DiMViGI framework to data at the participant-level and obtain distributions for the irreversibility of each tuple. As mentioned in the main manuscript, we analysed MEG recordings from 51 participants with 15 trials per participant. In the cohort-level analysis presented in the main manuscript, we constructed the in- and out-degree distributions using 51 *×* 15 = 765 samples of the multiplex network. In order to examine the spread between participants, we repeat the same analysis for each participant in isolation, using only the 15 associated trials. As a result the degree distributions are much more poorly estimated and produce much higher divergences. Nevertheless, we are able to quantify the irreversibility of each tuple of brain regions for each participant and examine the distribution.

Figure 1 shows the results of the DiMViGI analysis for the participant-level data distributions of the irreversibility for each tuple in each level. Panel a) shows the schematic representation of the 6 regions of interest (ROIs) that correspond to variables in the multivariate time-series. The icons on the *x*-axis of the subsequent panels indicate which ROIs are included in each tuple. Panels b-f) show the participant-level distributions for 1-5 order respectively. We run one-way ANOVAs and find that, at each level 1-5, the tuple is a significant predictor of irreversibility (*p <* 0.00001). In addition, we run paired *t*−tests to see which tuples at each level are significantly different in a pairwise comparison applying a Bonferroni correction for multiple comparisons at each level *k* [1]. Figure 2 displays the significance results of the pairwise *t*−tests. The corrected significance of each comparison is denoted as follows: (ns) if *p >* 0.05; (*) if *p <* 0.05; (**) if *p <* 0.01; (***) if *p <* 0.001 and (****) if *p <* 0.0001. Figure 2 shows that at level 1, the difference between each tuple in pairwise comparison is significant (*p <* 0.0001). In addition, it shows that at levels 2-5, there is a mixture of significant and not significant differences depending on the number of ROIs in common between the compared tuples.

Finally, we compare the participant-level analysis to the cohort-level analysis by calculating the ranking of tuples at each level for each participant and comparing it to the cohort-level ranking, using Spearman’s *ρ*. In addition, for each participant at each level, we calculate Pearson’s *r* (correlation coefficient) between the participant level and the cohort level. Panel a) of Fig 3 shows the *ρ* for each participant at each level when compared to the cohort-level ranking. Panel b) of Fig 3 shows the *r* for each participant at each level when compared to the cohort-level measurements. Both show that at lower orders (1-3), the measurements, and rankings, obtained from the participants in isolation agree closely with the cohort-level results. However, at higher order (4-5), the low number of samples, 15, in the participant-level analysis is not enough to accurately estimate the high-dimensional degree distributions leading to a lack of agreement between the well-estimated cohort analysis and the poorly-estimated participant analysis.

**FIG. 1.**
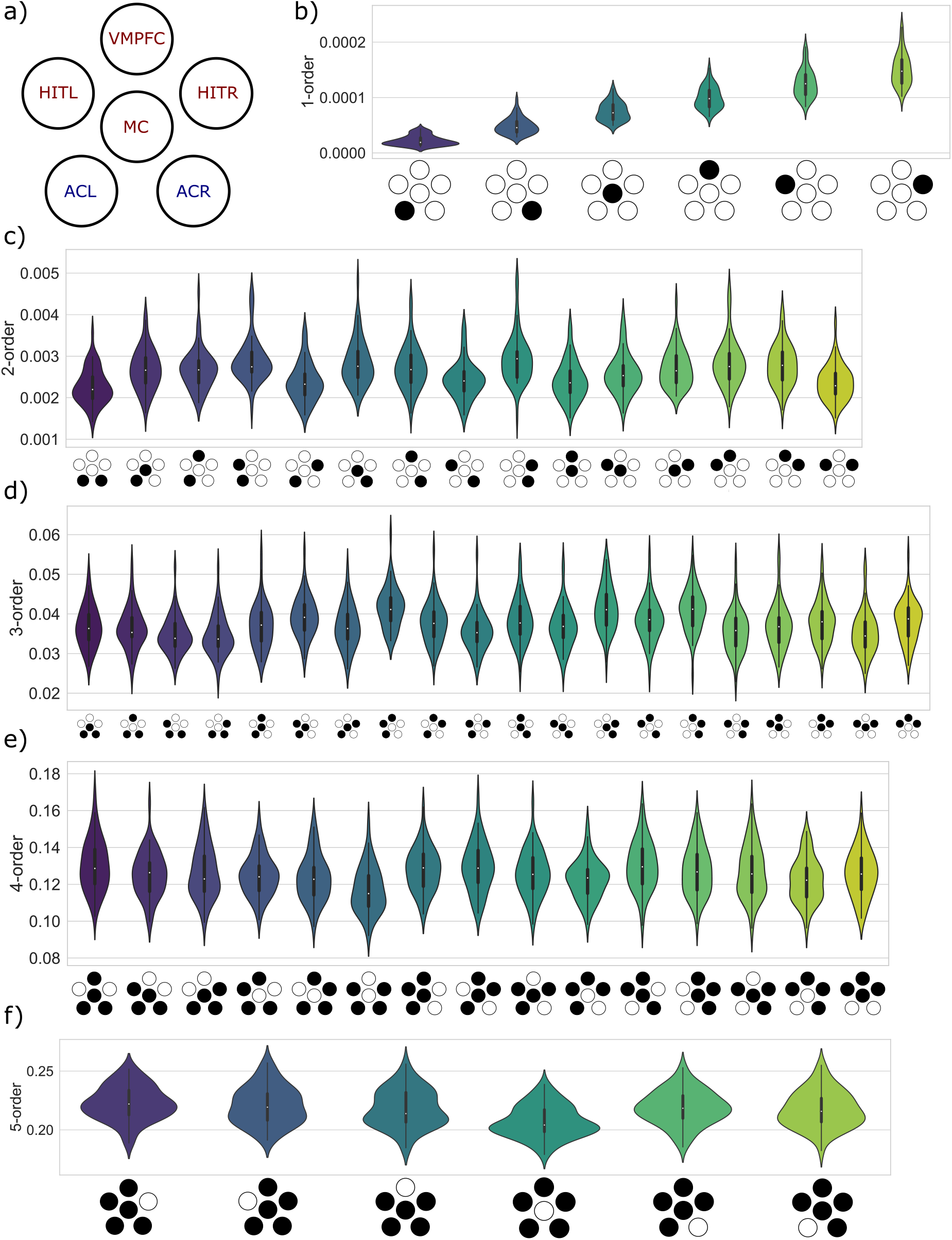
Participant distribution of irreversibility for each tuple. a) Schematic representation of the 6 brain regions of interest (ROIs) in the MEG recordings. The icon in the following panels indicates which regions are included in each tuple. b) 1-order irreversibility distribution for each ROI in isolation. The results follow the same hierarchy as the cohort-level analysis in the main manuscript. c) 2-order irreversibility distribution for each pairs of ROIs. d) 3-order irreversibility distribution for each triplet of ROIs. e) 4-order irreversibility distribution for each quadruplet of ROIs. f) 5-order irreversibility distribution for each quintuplet of ROIs.

**FIG. 2.**
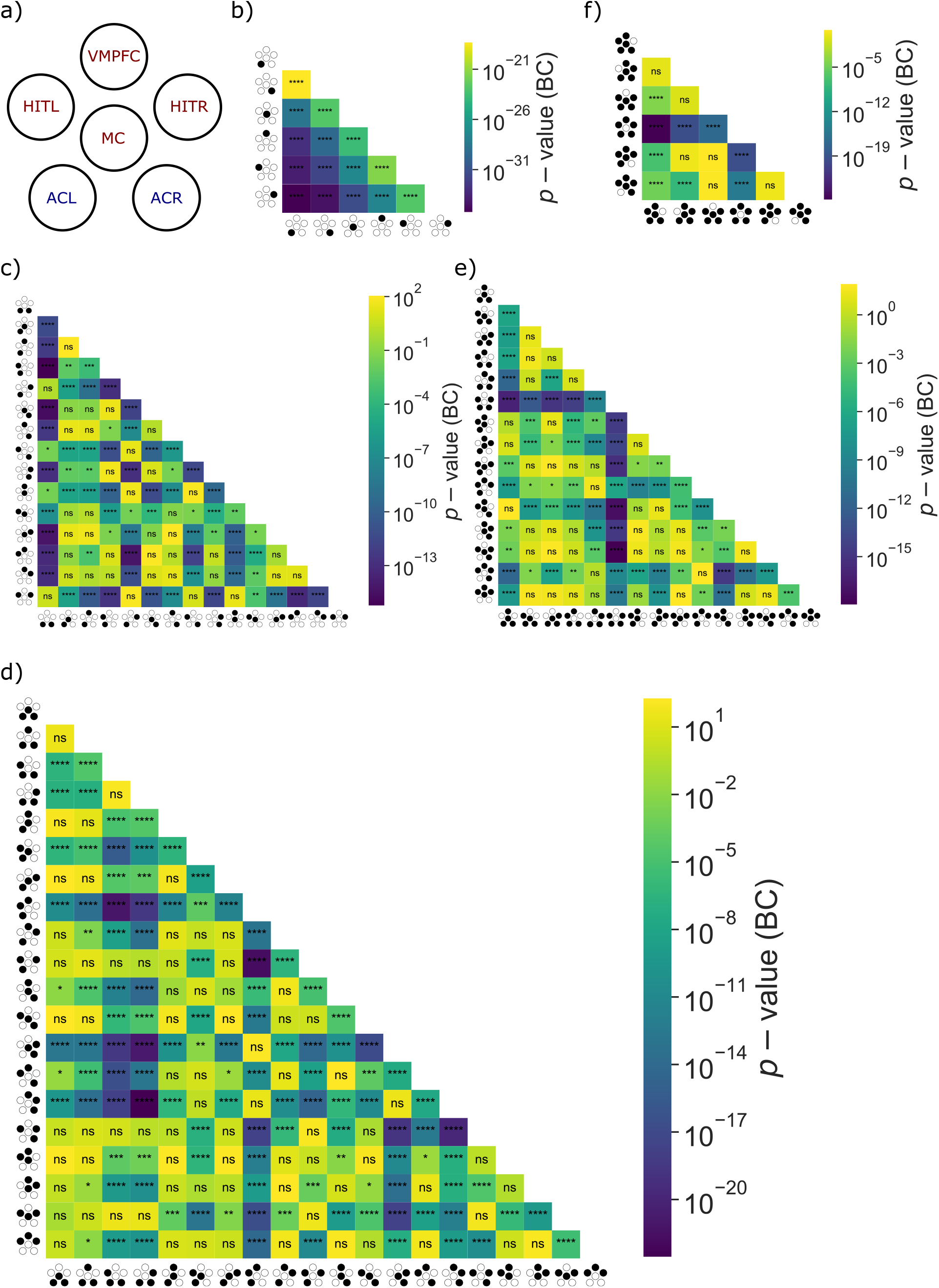
Results of pairwise *t*−tests (paired) between all pairs of *k*-tuples at each order *k* with a Bonferroni correction for multiple comparisons [1]. The corrected significance of each comparison is denoted as follows: (ns) if *p >* 0.05; (*) if *p <* 0.05; (**) if *p <* 0.01; (***) if *p <* 0.001 and (****) if *p <* 0.0001. Panel a) shows a schematic representation of the 6 brain regions of interest (ROIs) in the MEG recordings. The remaining panels show the results for b) singletons, c) pairs, d) triplets, e) quadruplets, f) quintuplets. There is a mixture of significant and not significant differences depending on the number of ROIs in common between the compared tuples

**FIG. 3.**
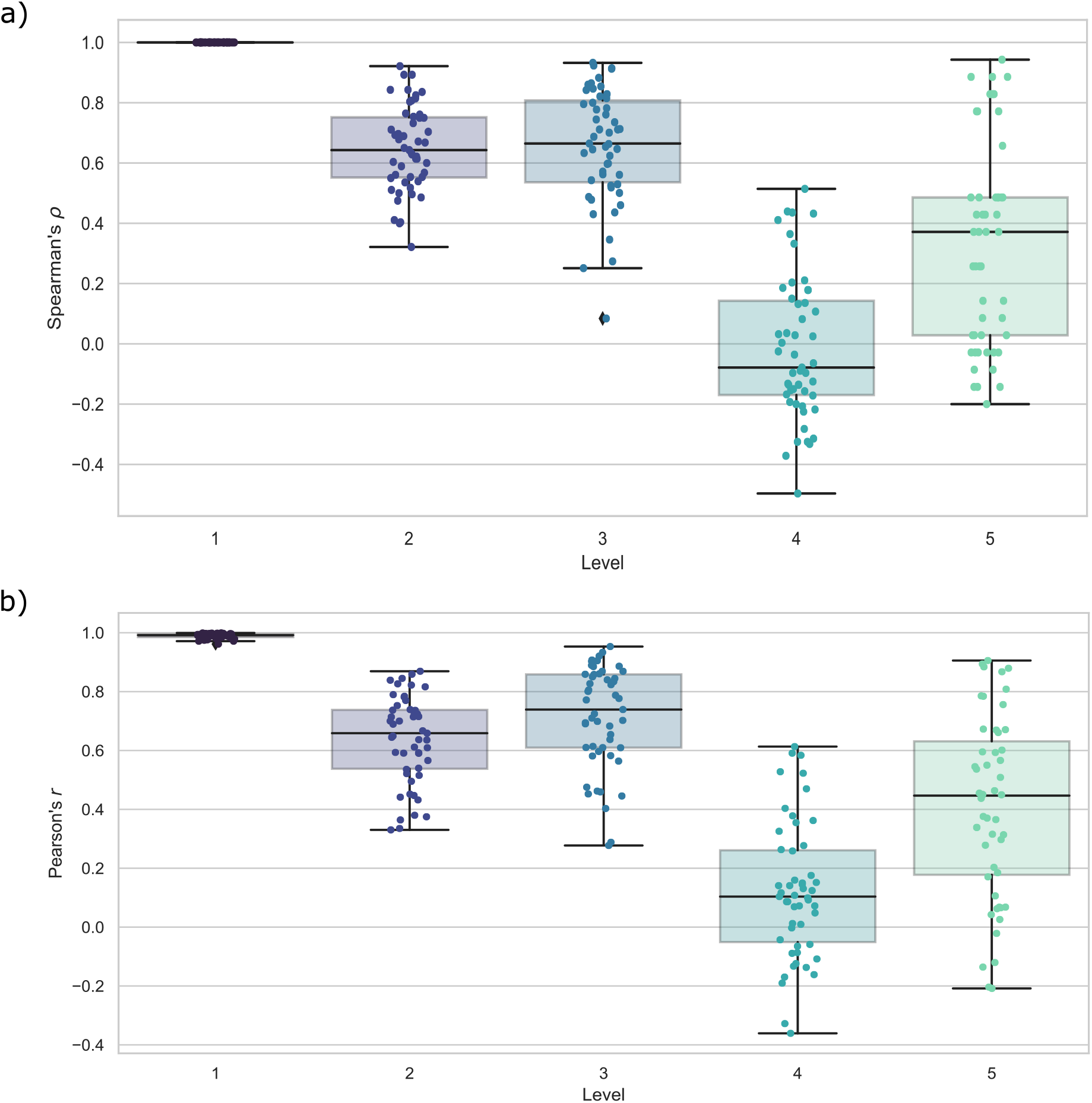
Correlations between participant and cohort level results. a) Spearman’s *ρ* coefficient for the ranking of tuples at each level for each participant when compared to the ranking obtained from the cohort-level analysis. b) Pearson’s *r* correlation coefficient for the measurements of tuples at each level for each participant when compared to the measurements obtained from the cohort-level analysis. The figure shows that at lower orders, the participant distributions agree more closely with the cohort-level analysis. However, at higher orders, the degree distributions are more poorly estimated leading to low agreement between the cohort and participant-level analysis.

### III. VALIDATION OF RESULTS AGAINST SURROGATE DATA FROM SHUFFLING TIME-SERIES

When measuring information-theoretic quantities from finite-length series, a bias is introduced. Finite-length time-series have a level of irreversibility that emerges from their finiteness as opposed to the temporal organisation of the data [2, 3]. In order to validate that the measured irreversibilities are significant and therefore emerge from the temporal structure of the data, we must compare them to surrogate data. When generating surrogate data, we aim to break the temporal correlations and restore detailed balance. In order to do this, we randomly shuffled the time-series in time. This means that the number of occurrences of each state remains the same as the original data, but the sequence of states is now randomised thereby restoring detailed balance, as shown by Lynn et al [3]. The irreversibility measured in the shuffled time-series is an estimate of the ‘noise-floor’ which stems from finiteness of the series. If the irreversibility of the true signal is significantly higher than the noise floor, then this is due to the system violating detailed balance in the underlying dynamics.

Other common approaches for generating surrogate time-series such as phase randomisation or Fourier transform surrogates preserve the temporal structure of the time-series and are therefore unsuitable for this application [4, 5].

Figure 4 shows the comparison of the measurements in the original MEG time-series and its randomly shuffled surrogate. For each tuplet the difference between the shuffled and original time-series is significant (*p <* 0.0001). This shows that the irreversibility measured using the DiMViGI framework is a significant statistical feature of the multivariate time-series as the shuffled data is measured to be far more reversible using the DiMViGI framework.

### IV. ESTIMATING FINITE-DATA ERRORS USING SUB-SAMPLING

In addition to shuffling the time-series, another approach for estimating errors that arise from finite-data, is to employ a sub-sampling approach [2, 3, 6]. As the DiMViGI framework, when applied to all participants, calculates a single quantity for the irreversibility of each tuple, we need to estimate the size of the error as well as how this error evolves with the amount of data that is available. To do this, we consider the relationship between irreversibility and the *inverse-data-fraction* (IDF) which is 1 divided by the fraction of the data that is used to estimate the irreversibility. We do so by randomly sampling (without replacement) 12 and then 9 trials per participant in a hierarchical fashion, meaning 9 are chosen from the 12 which are chosen from the 15. Whilst keeping the contribution of each participant equal, this allows us to calculate the irreversibility using fractions 1, 0.8, 0.6 and of the full dataset corresponding to IDF=1, 1.25 and 1.67 respectively. Importantly, there are many ways to sub-sample the data which allows one to obtain a number of different estimates, and thus an uncertainty measure, for the irreversibility at IDF=1.25 and 1.67. In the following analysis, we use 20 random sub-samplings. Some studies in neural spike trains linearly extrapolate this to the *infinite data limit*, corresponding to IDF=0 [6–10]. However, we find that the relationship between irreversibility under DiMViGI and IDF is nonlinear. Therefore this extrapolation may yield nonphysical (negative) values of irreversibility and thus is not suited for our analysis. We note that such analysis is often useful when attempting to measure the *true* value of a quantity in particular units, for comparison to other experiments. In our study, the measurements obtained by DiMViGI are relative quantities, to be compared within a given level *k*.

Figure S5 shows the results of sub-sampling at order 1. Panels a) and c) show that the 1-order irreversibilities measured at IDF = 1.25 and 1.67 agree closely with IDF =1, the results in the main manuscript. Furthemore, panels b) and d) show the pairwise comparisons between the distributions of irreversibility measured for each ROI at both IDF= 1.25 and 1.67 (paired *t*−test, BC). We find each comparison is significant (****, *p <* 0.0001) indicating that the differences between ROIs are much larger than the errors in the measured irreversibility. Figures S6, S7 show similar results at order 2, pairs of regions, and order 3, triplets, where pairwise comparison shows a range of significant differences. Figures S8 & S9 show that the results for orders 4 & 5, quadruplets and quintuplets, agree between IDF=1 and IDF=1.25, 1.67. Furthermore, we find that most quadruplets and all quintuplets have significantly different levels of irreversibility. These results suggest that biases and errors in the irreversibility measurements are typically smaller than differences in measurements between tuples and further validates our approach of identifying particularly (ir)reversible tuples. We note that the quintuplet analysis for the sub-sampling uses degree-limiting with a maximum degree of 45, which, as shown in Section 8, minimally affects the results whilst improving the computational efficiency.

### V. THE DIMVIGI FRAMEWORK FACTORISES FOR INDEPENDENT VARIABLES

To illustrate that the DiMViGI framework indeed can differentiate higher order interactions from the composition of lower order ones, we consider the framework applied to a *k*-tuple of variables, (*x*_1_, …*x*_*k*_). First we assume that *x*_1_ is independent of (*x*_2_, …, *x*_*k*_) and show that we can write the irreversibility of the *k*−tuple as the sum of the irreversibility of *x*_1_ plus the irreversibility of the (*k*− 1)−tuple (*x*_2_, …, *x*_*k*_). Inductively, we can show that the DiMViGI framework factorises for independent variables meaning that the irreversibility of their interaction is merely the sum of the irreversibility of each variable in isolation. This validates that the framework is truly capturing multilevel irreversibility.

**FIG. 4.**
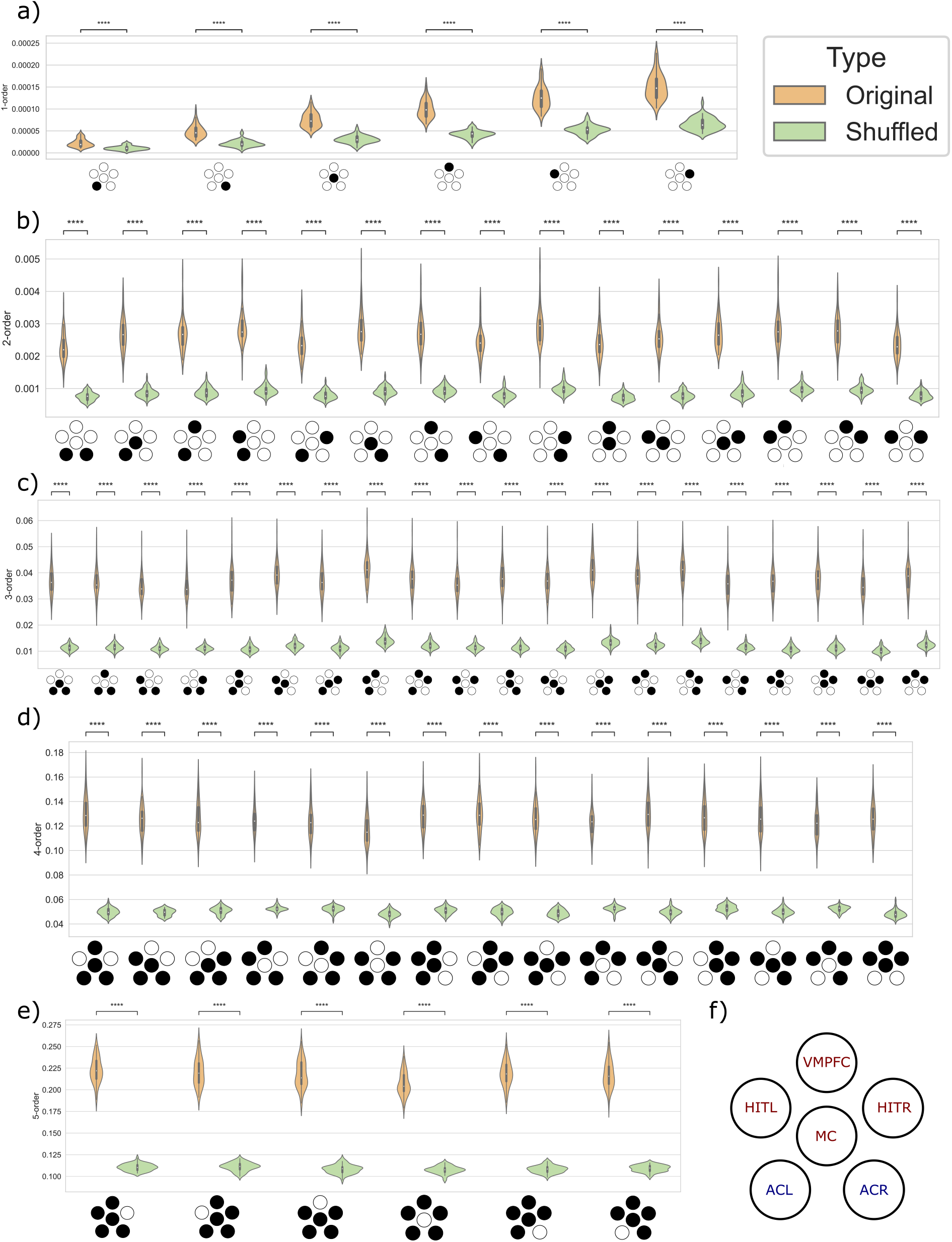
Comparison of original neural recording against surrogate data obtained via shuffling the time-series in time. The difference between the original and shuffled data for each tuple at each level is significant. We label (****) if *p <* 0.0001. a) 1-order irreversibility is significant (*p <* 0.0001) for each ROI when compared to shuffled data. b) 2-order irreversibility is significant (*p <* 0.0001) for each pair of ROIs when compared to shuffled data. c) 3-order irreversibility is significant (*p <* 0.0001) for each triplet of ROIs when compared to shuffled data. d) 4-order irreversibility is significant (*p <* 0.0001) for each quadruplet of ROIs when compared to shuffled data. e) 5-order irreversibility is significant (*p <* 0.0001) for each quintuplet of ROIs when compared to shuffled data. f) Schematic representation of the 6 brain regions of interest (ROIs) in the MEG recordings. The icon in the preceding panels indicates which regions are included in each tuple.

Consider (*x*_1_, …, *x*_*k*_) such that *x*_1_ is independent of the other variables. As *x*_1_ is independent, the edges in the associated layer of the multiplex network are also independent. As a result, the joint (in- and out-) degree distribution of the multiplex factorises as follows,

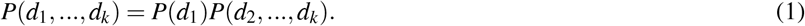

Under the DiMViGI framework, we quantify the irreversibility of the tuplet as

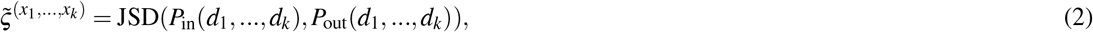

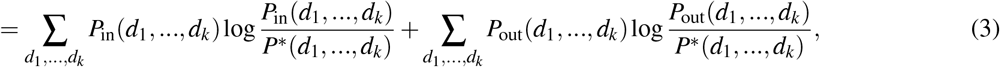

where 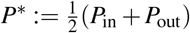 We focus first on the term concerning the in-degree distribution and use the independence of *x*_1_ and the properties of logarithms to factorise and simplify this expression,

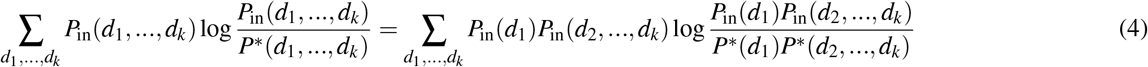

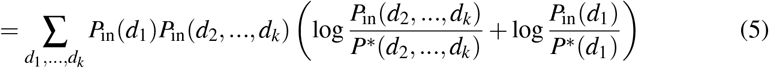

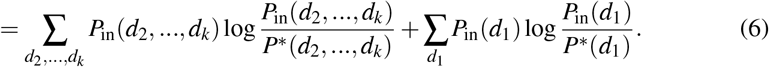

By symmetry the same is true for the term concerning the out-degree distribution. Substituting in the simplified expression, we get

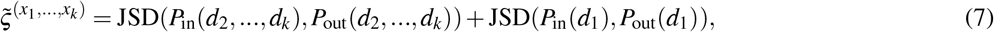

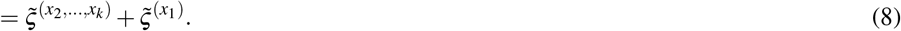

This indicates that in this *k*-dimensional system that does not contain a genuine *k*-order interaction, the irreversibility of the *k*−tuple simply decomposes into the sum of non-independent tuples. By induction, for a *k*−tuple where all variables are independent, the irreversibility fully decomposes into the sum of the 1-order irreversibilities,

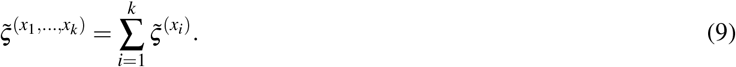

### VI. UNIQUE CONTRIBUTIONS FROM HIGHER ORDER INTERACTIONS

In our analysis, we have considered the irreversibility of multilevel interactions. However, for a given *k*−order interaction, we have measured the irreversibility of the combined *k*−tuple. This is in contrast with the ‘unique’ irreversibility that is contributed purely by the *k*-body interaction, discounting the *j*−body interactions for *j < k* that are included within this *k*−tuple.

Within the theory of higher order interactions, this distinction represents the difference between a hyper-graphical structure and a simplicial complex [11]. In the former, a *k*-body interaction does not comprise of lower-order components, whereas in the latter, every lower order relationship must exist to define a higher order one i.e. a 3−order triangle relationship requires all the edges of the triangle to be included.

Within the lens of irreversibility, we note that the decomposition proposed by Lynn et al [6, 12] specifically considers the unique contributions to the global irreversibility. Alternatively, in the manuscript, we present a method that captures the irreversibility of path projected into the portion of state-space defined by a tuple,

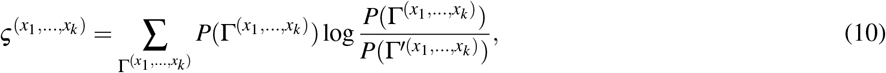

which is not equivalent to the unique contribution. In this section, we relate our framework more closely, but still not equivalently, to the decomposition in Ref. [12] by measuring the unique contribution of the *k*-body interaction to 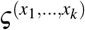 We do this by recursively subtracting the irreversibility of sub-tuples Ω ⊂ {*x*_1_, …, *x*_*k*_}, from the quantity 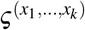 In such a way we define the unique contribution to the 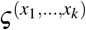 of the *k*−body interaction (*x*_1_, …, *x*_*k*_) as,

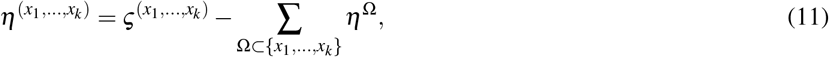

which is calculable by noting that 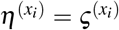 We are able to show that, using this framework, the results are highly correlated, indicating that higher order interactions dominate the irreversibility in these large-scale neural recordings. This stands in contrast with results obtained in spike-train data that indicate that, at the neuronal level, pairwise interactions dominate [6, 12].

We note that this approach captures the unique contributions of *k*-body interactions by considering the following

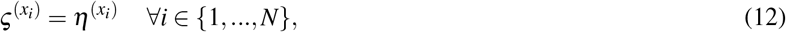

i.e. the combined and unique irreversibility at 1-order is equivalent. Next we note that for two independent variables, the irreversibility factorises, under the DiMViGI framework,

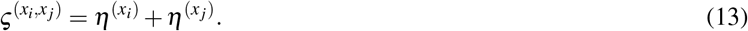

For independent variables *x*_*i*_, *x* _*j*_, we expect 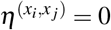. Therefore, it is natural to define,

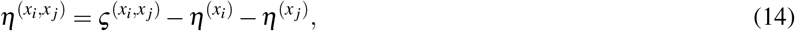

which is positive for correlated variables and vanishes for independent variables. By definition, it captures the irreversibility of the pairwise interaction, discounting the singleton dynamics. In this fashion, we can recursively calculate the unique contributions at *k*-order using the unique contributions at *j*-order for 1 ≤ *j < k*. Concretely, we have,

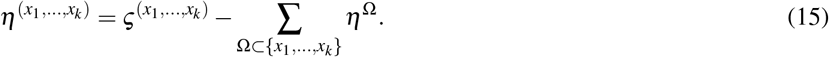

Figure 10 shows the contrast between the unique and combined irreversibilities for tuples at levels *k* = 2, 3, 4 at the cohort-level. We do not consider 1-order as the unique and combined values are equivalent. Furthermore, we cannot consider *k* = 5 as we employ degree-limiting (see Section VIII) for computational efficiency at this level. As a result, we consciously underestimate the irreversibility at 5-order which leads to negative values when inputting these measurements into equation 15. Panel a) of Fig 10 shows a small level of contrast between the unique and combined pairwise dynamics. This indicates that the irreversibility of pairwise interactions dominates the irreversibility of singleton dynamics. Furthermore the general hierarchy is preserved. Panels b-c) show similar results with increasing levels of contrast. However, this increasing contrast is due to the combinatorics of higher order interactions. In particular, as we increase the level *k*, we are subtracting more terms when isolating the unique contribution. However, this difference is overstated, as panel d) shows that the correlation between unique and combined measurements is almost perfect. This indicates that at a given level *k*, the *k*-body interaction dominates the lower level interactions and contributes the most to the irreversibility. This result is both a consequence of the method, and the spatially-coarse, low-dimensional data under consideration. It further suggests that, whilst the DiMViGI framework can be used to compare irreversibility between levels, it is most useful for comparing tuples within a given level.

### VII. VALIDATION USING SIMULATED DATA FROM THE MULTIVARIATE ORNSTEIN-UHLENBECK PROCESS

Next, we aim to validate our technique against simulated time-series. We choose the multivariate Ornstein-Uhlenbeck as this is one of the few models that has a known rate of entropy production [13]. Furthermore, this model has been fit to neural recordings in the past in order to estimate the entropy production rate [14].

#### A. Multivariate Ornstein-Uhlenbeck process

The Orstein-Uhlenbeck process models the velocity of a particle in Brownian motion [15]. In its generalised multivariate form, we consider *N* particles with coupled stochastic dynamics given by the equation,

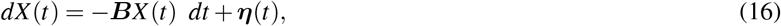

where *X* (*t*) ∈ ℝ^*N*^. The friction −***B*** ∈ ℝ^*N×N*^ is a stable matrix, meaning that every eigenvalue has strictly negative real part. The additive noise, ***η***(*t*), is Gaussian and has covariance ***D*** ∈ ℝ^*N×N*^,

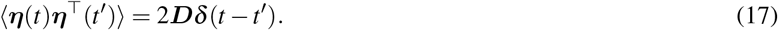

***D*** is a symmetric, positive definite matrix, and so we can calculate ***L***, its Cholesky decomposition, where ***L*** satisfies ***D*** = ***LL***^⊤^and ***L*** is lower triangular [13]. As a result, we can write the system as a Langevin equation,

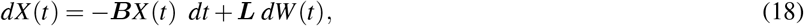

where *W* (*t*) represents a *N*-dimensional Wiener process with independent components. The individual trajectories of a mOU are always reversible, yet at the macroscopic level, irreversibility can emerge. The macroscopic process is known to be reversible if ***BD*** is symmetric i.e.,

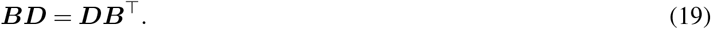

Note that ***D*** is always symmetric, whereas, in general, ***B*** is not. Furthermore, the covariance, ***S***, of the stationary state can be defined implicitly in terms of ***B*** and ***D*** by the Lyapunov equation,

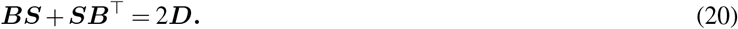

In the case that the process is reversible, we can use the criterion (19) to write ***S*** explicitly,

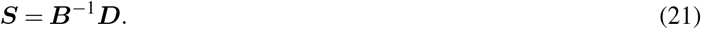

In the case that the process is irreversible, obtaining an explicit form for ***S*** is not as simple. Instead, we parameterise the level of asymmetry using the Onsager matrix of kinetic coefficients and a matrix ***Q***, that represents the asymmetry,

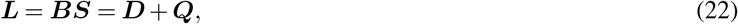

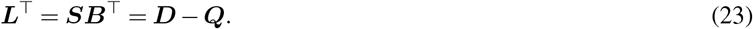

As shown in [13], the entropy production rate for the multivariate Ornstein-Uhlenbeck process can be written in terms of the matrices ***B, D*** and ***Q***. The rate of entropy production is given by,

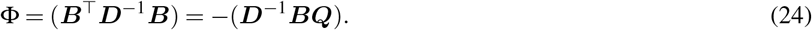

Clearly, when the process is reversible, ***Q*** = 0 and thus Φ = 0. In general, the matrices ***S*** and ***Q*** cannot be determined in closed form and so Φ does not have a closed form expression. However, in the case *N* = 2 or in the presence of appropriate symmetries in the matrices ***B*** and ***D***, a closed form expression can be derived for Φ [13].

##### 1. *The case N* = 2

In the case *N* = 2, the Lyapunov equation (20) has a closed form solution, and therefore the entropy production rate can be explicitly expressed as a function of the entries of ***B*** and ***D*** [13].

Consider the matrices,

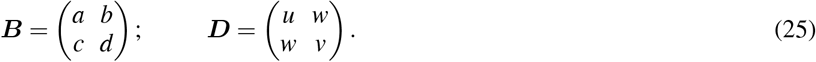

In this case, the rate of entropy production is given explicitly by the formula,

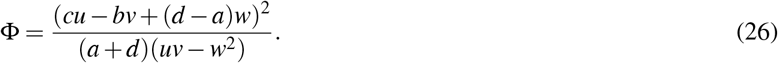

Clearly we have Φ = 0 if and only if the reversibility criterion,

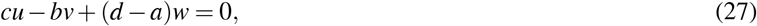

is satisfied [13].

##### 2. Cyclic symmetry

Consider the situation where the variables live on a ring with *N* sites where the dynamics are invariant to translations of the ring. This results in the matrices ***B*** and ***D*** being circulant, i.e.

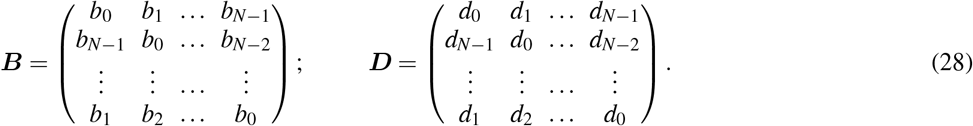

As ***D*** is assumed to be symmetric, this imposes the additional restriction that *d*_*N*−*i*_ = *d*_*i*_. In this case, the rate of entropy production has a closed form expression,

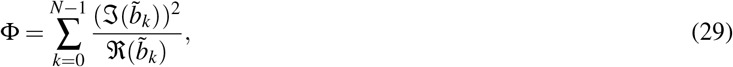

where 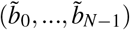 is the discrete Fourier transform of the vector (*b*_0_, …, *b*_*N*−1_), ℑ(*·*) represents the imaginary part of a number and ℜ(*·*) represents the real part [13]. Recall that for a circulant matrix, the Fourier modes of (*b*_0_, …, *b*_*N*−1_) coincide with the eigenvectors of ***B***.

#### B. Example processes validating the DiMViGI framework

Using the cases where we can calculate the explicit rate of entropy production, such as those detailed above, we can construct example processes and compare the measurements from our technique to the global rate of irreversibility. Figure 11 shows the results of these numerical experiments.

##### 1. Example 1

We first consider Example 1, a 2-dimensional process with friction and noise given by,

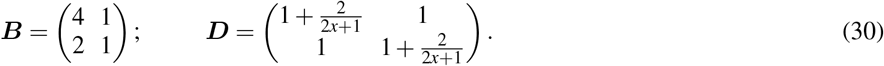

This gives rate of entropy production,

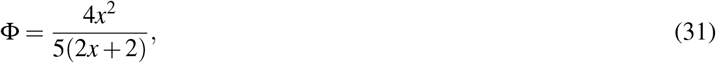

which vanishes for *x* = 0, corresponding to a reversible process. Furthermore, as *x* increases from 0, the rate of entropy production grows linearly with *x*.

We numerically sample paths from this process for values *x* = 0, 0.5, 1, …, 10 using an Euler-Maruyama scheme. We sample paths of length *T* = 500 with a time-step of Δ*t* = 0.01 and keep only the last 2000 time-steps of the process, to avoid boundary effects.

As shown in Panel a) Fig. 11, we can see that the first order irreversibility captured by the DiMViGI techniques shows no correlation with the global rate of entropy production. This is because individual trajectories of the mOU are reversible. As a result, what is plotted is numerical error associated with finite trajectories which has no correlation with the parameters or Φ. On the other hand, the second order irreversibility of the pair (*x*_1_, *x*_2_) is capturing the global rate of entropy production as this interaction produces all the entropy in the system. As a result we can see the strong linear correlation between the 2-order irreversibility and Φ.

##### 2. Example 2

We consider Example 2, a circulant 3-dimensional process with a strong triplet interaction,

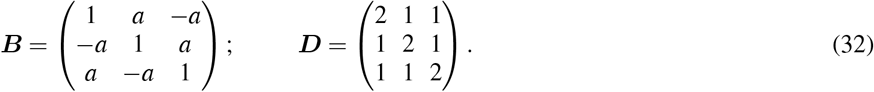

This gives rate of entropy production,

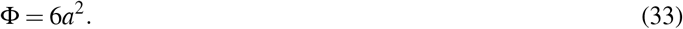

For *a* = 0.5, 1, …, 8.5, we sample paths with the same methods as before and estimate the irreversibility of the interactions in the system.

As shown in Panel b) of Fig. 11, the individual trajectories are again uncorrelated with the global rate as they are reversible. Both the pair and triplet dynamics are strongly correlated with the global rate of entropy production. Whilst we do not know how much each pair contributes to Φ, the circular symmetry of the process suggests the dynamics of pairs should be identical, which we see here.

##### 3. Example 3

We consider Example 3, a circular 3-dimensional process with only pairwise drift interactions, given by,

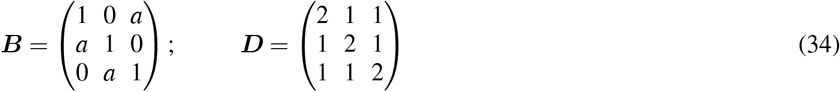

which, again, gives rate of entropy production,

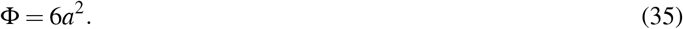

Paths are sampled for value *a* = 0.1, 0.2, …, 1.8. Whilst each component is only coupled to itself and one other component directly in the drift matrix, it is coupled to the entire system via the noise matrix and indirectly via the dynamics of the other components. For example, even though *x*_2_ does not appear in the drift term for *x*_1_, they are correlated through shared noise and via *x*_3_. For this reason, the difference between Example 2 and Example 3 is not extreme. As can be seen in Panel c) of Fig. 11, we get almost identical dynamics of the measure. Whilst, we aim to distinguish between Example 2 and Example 3, by restricting to pairwise or triplet dynamics, we note that the mOU is a linear system that can be decomposed into its pairwise interactions, meaning it cannot produce genuine higher order effects [11]. However, we are restricted in this analysis to this model as the explicit entropy production rate is known.

##### 4. Example 4

We consider Example 4 which is 4-dimensional and non-circulant. As a result, we no longer have the exact solution for the entropy production rate and must estimate this quantity numerically. Example 4 has drift and covariance,

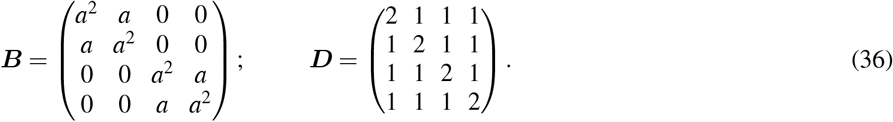

This system is interacting as a 4-dimensional system as it is coupled through the noise dynamics. However, in the drift term we have two subsystems where (*x*_1_, *x*_2_) interact strongly as do (*x*_3_, *x*_4_), but these pairs are drift-wise disjoint. In order to numerically estimate the entropy production rate, we estimate the covariance matrix from the sampled paths,

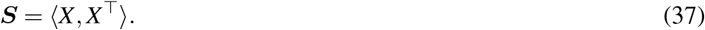

Next, we can calculate the asymmetric part of the Onsager matrix,

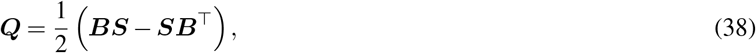

which can be used to calculate the entropy production rate,

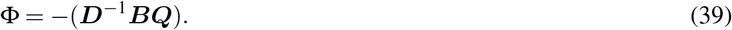

We sample paths for values *a* = 2.5, 2.7,…, 4.9, but we do not know how Φ scales with *a*. Panel d) of Fig. 11 is harder to interpret than for the previous examples, as the numerical approach produces greater variance in the plot. However, we plot least-square regression lines for each tuple. At 1-order, the lines are almost flat, as expected as there should be no correlation between the reversible individual trajectories and the underlying rate of entropy production. At 2-order, we have the most important result, which is that whilst the reversibility of all pairs scales with entropy production, the strongly interacting pairs (*x*_1_, *x*_2_) and (*x*_3_, *x*_4_), the upper two lines, are more irreversible. At 3-order, all the interactions produce almost identical amounts of irreversibility, which is to be expected as each triplet contains a strongly interacting pair and one component from the other pair, leading to a symmetry in the dynamics. Finally, the irreversibly of the quadruplet, the entire system, scales linearly with the underlying entropy production rate.

We note that the mOU is not a truly higher-order system as it is linear and the interactions can be seen as pairwise, but we are restricted to this model as it has a known rate of entropy production and producing continuous dynamics. Other techniques have validated their techniques on chaotic processes [16] or symbolic dynamics i.e. Ising model [3]. However, deterministic chaos and thermodynamic irreversibility are not equivalent. Furthermore, these processes do not allow one to scale the number of variables, nor the level of irreversibility, arbitrarily. On the other hand, by varying the thermodynamic temperature, one can vary the irreversibility of the Ising model, but the visibility graph is designed to capture correlations in continuous rather than binary series yielding this model unsuitable. For this reason, we opt exclusively for the mOU as studied here.

### VIII. VARYING THE MAXIMUM DEGREE IN THE SUPPORT OF THE DEGREE DISTRIBUTIONS

The DiMViGI framework projects the high-dimensional, continuous state-space of the multivariate time-series into a discrete and low-dimensional representation using the visibility graph, thus reducing the computational cost of calculating information-theoretic quantities [17, 18]. However, the combinatorial complexity of considering every possible tuple in a system can be restrictive. Furthermore, estimating high-dimensional degree distributions can also be computationally demanding in terms of computer memory. A simple method for improving the memory efficiency of the DiMViGI framework is to cap the maximum degree in the support of the degree distribution. The degree distribution of the visibility graph typically decays exponentially as the degree increases [17, 19, 20]. As a result, when limiting the degree, we are removing minimal information. Moreover, a *k*-dimensional distribution with maximum degree *d*_max_ contains 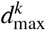 entries. Therefore, degree-limiting has an exponential reduction in the memory usage of the DiMViGI implementation. In our analysis presented in the main manuscript, we employed degree limiting in the case of *k* = 5, where we enforced *d*_max_ = 75. In this section, we present a systematic analysis of the effect of degree limiting for each tuple at each level. We implement this limiting by enforcing that if a node has a degree greater than 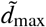, we set its degree to 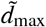 in the distribution.

First, we note that, in practice, the restriction causes us to underestimate the irreversibility of the tuple. However, this is not mathematically guaranteed. For a tuple, (*x*_1_, …, *x*_*k*_), we denote the irreversibility with full support to be 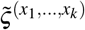 and the irreversibility with limited support to be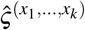. Therefore, the difference is,

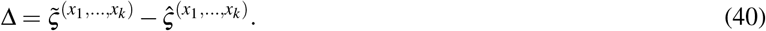

The sign of Δ reflects whether we are over or underestimating the irreversibility using the limited support. We recall the definition of JSD between distributions *P* and *Q*,

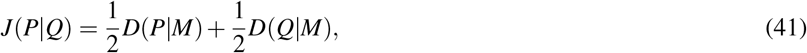

where 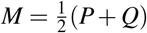 is an averaged distribution and *D*(*·*) represents the KLD, given by,

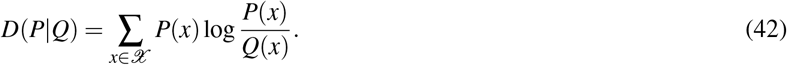

Therefore,

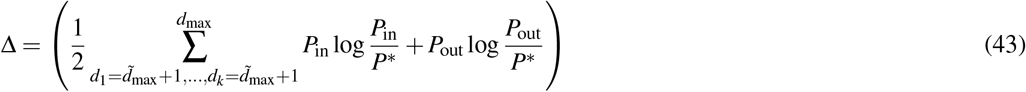

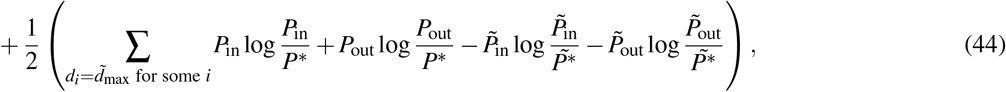

where *P*_in_, *P*_out_ are the in- and out-degree distributions with the full support; 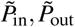 are the in- and out-degree distributions with the limited support and *P*^*^, 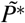 are the averaged in-out distributions. In other words, the limited and full degree distributions overlap for all degrees 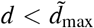 and therefore cancel when we take the difference. By truncating the distribution we are neglecting a number of positive terms from the full support. However, we must also consider that these edges have not been simply deleted, but are now included in overestimating the probability that a node has degree 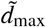, hence *P* and 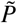 differ when a node has degree 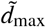 in some layer. We can rewrite Δ as,

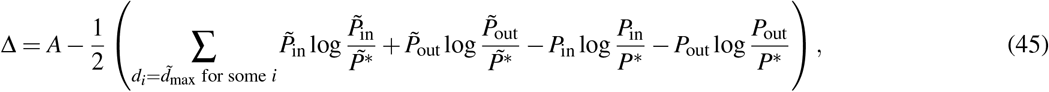

where *A* ≥ 0. Therefore, if the subtracted sum is less than *A*, we will underestimate the irreversibility, but if it greater than *A* we will overestimate the irreversibility. As shown in Figure 12, in practice, we consistently underestimate the irreversibility by limiting the degree, indicating that *A* is much larger than the subtracted term.

Figure 12 shows a systematic variation of the maximum degree. We perform the analysis with 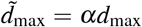 and *α* ∈ [0, 1], where *d*_max_ is the maximum degree of the full multiplex visibility graph (MVG) from the data for *k* = 1, …, 4 and *d*_max_ = 75 for *k* = 5. We vary *α* = 0, 0.1,…, 0.9, 1 and calculate the irreversibility of each tuple using the DiMViGI framework but with restricted degree distribution. In addition, we also show, for each value of *α*, the Pearson correlation coefficient, *r*, and Spearman’s rank, *ρ*, between the limited and full support values at each level. Panels a-e) show the effect on the irreversibilities of *k*-tuples with *k* = 1, …, 5 respectively and panel f) recalls the schematic representation of the regions of interest. For each level, the irreversibility monotonically increases as we increase *α*, confirming that degree-limiting underestimates the irreversibility. For lower orders (1-2), we see that the increase is linear. In particular, for the pairwise results, to get a strong correlation with the full support irreverisibility, one needs to use a large proportion of the *d*_max_. On the other hand, for the higher orders (3-5), the increase is sigmoidal. Panels c-e) indicate that even limiting to half of the maximum degree is sufficient for an almost perfect correlation with the original results. For order 4-5, we see that this also captures approximately 90% of the irreversibility. With an exponentially smaller distribution, one can capture almost equivalent information. This analysis indicates that degree limiting is a very practical and useful tool to maximise the memory efficiency of the DiMViGI framework at higher orders.

### IX. FORMULATIONS OF THE ENTROPY PRODUCTION RATE FOR MARKOVIAN AND NON-MARKOVIAN SYSTEMS

For certain systems in a stationary state, the entropy production rate (EPR) can be explicitly formulated.

In the case of discrete time, discrete space, Markovian dynamics, the entropy production rate simplifies to,

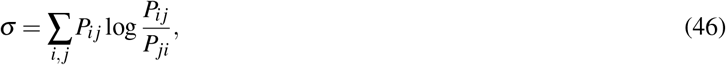

where *P*_*i j*_ is the join transition probability, *P*(*x*_*t*+1_ = *j, x*_*t*_ = *i*) [3].

For *l*−order Markovian dynamics, the rate of entropy production is given by,

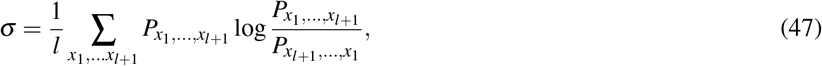

where 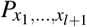 is the probability of observing the exact sequence of states 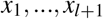[3].

For a general system with discrete states in discrete time, the rate of entropy production of a single trajectory is given by,

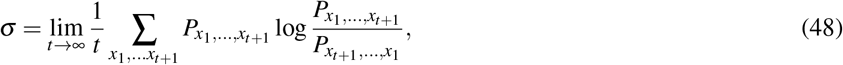

where 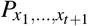 is the probability of observing the exact sequence of states 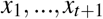 [3].

For continuous time Markovian dynamics, the time-dependent rate of entropy production is given by,

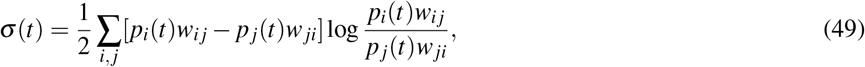

where *p*_*i*_*(t)* is the instantaneous probability distribution and *w*_*i j*_ are the transition rates [21].

For Markovian dynamics in continuous space and time, given by the Langevin equation,

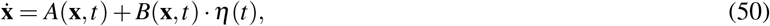

we can write an equivalent Fokker-Planck equation,

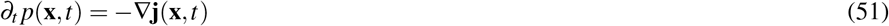

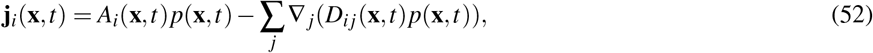

where,

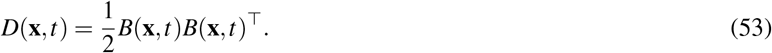

Following [22], the time-dependent entropy production rate is given by,

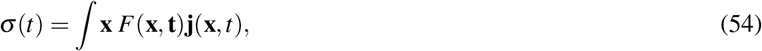

where,

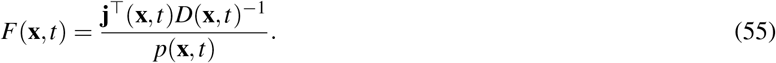

### X. EXPERIMENTAL PARADIGM AND MEG RECORDINGS

In this section, we provide additional information about the experimental paradigm, acquisition and pre-processing of the MEG recordings.

#### A. Experimental paradigm

We employed an old/new paradigm auditory recognition task [23–28]. Participants listened the first four bars of the right-hand part of Johann Sebastian Bach’s Prelude No. 2 in C Minor, BWV 847, twice and were asked to memorise it to the best of their ability. Next, participants listened to 135 five-tone musical sequences, corresponding to 27 trials in 5 experimental conditions, of 1750 ms each and were requested to indicate if the sequence belonged to the original music or was a variation. The experimental conditions corresponded to systematic variations on the position of the first varied tone in the sequence. For a detailed description and analysis of the different experimental conditions, see Bonetti et al [23]. We consider one experimental condition, where participants recognised the original, previously memorised sequences.

#### B. Data acquisition

The MEG recordings were taken in a magnetically shielded room at Aarhus Univeristy Hospital (AUH), Aarhus, Denmark on an Elekta Neuromag TRIUX MEG scanner with 306 channels (Elekta Neuromag, Helsinki, Finland). The sampling rate was 1000 Hz with analogue filtering of 0.1-330 Hz. Before taking the recordings, we registered the head shape of participants and the position of four Head Position Indicator (HPI) coils with respect to three anatomical landmarks using a 3D digitiser (Polhemus Fastrak, Colchester, VT, USA). We used this recording to co-register MRI scans with the MEG recordings. During the MEG recordings, the HPI coils continuously registered the localisation of the participant’s head which was then used for movement correction. Furthermore, heartbeats and eye-blinks were recorded with two sets of bipolar electrodes which were then used, further along the pre-processing pipeline, to remove artefacts from the MEG recordings. The MRI scans were taken on a CE-approved 3T Siemens MRI-scanner at AUH. The MRI data consisted of structural T1 (mprage with fat saturation) with a spatial resolution of 1.0 × 1.0 × 1.0 mm and the following sequence parameters: echo time (TE) = 2.61 ms, repetition time (TR) = 2300 ms, reconstructed matrix size = 256 × 256, echo spacing = 7.6 ms, bandwidth = 290 Hz/Px. The MRI and MEG recordings were acquired on two separate days.

#### C. MEG data pre-processing

Firstly, the raw MEG sensor data (204 planar gradiometers and 102 magnetometers) was pre-processed using MaxFilter to attenuate external interferences [29]. Next, signal space separation was applied with MaxFilter parameters: spatiotemporal signal space separation [SSS], down-sample from 1000Hz to 250Hz, movement compensation using cHPI coils [default step size: 10 ms], correlation limit between inner and outer subspaces used to reject overlapping intersecting inner/outer signals during spatiotemporal SSS: 0.98). Then the data was converted into Statistical Parametric Mapping (SPM) formatting and further pre-processed in MATLAB (MathWorks, Natick, MA, USA) using in-house-built codes (LBPD, available at https://github.com/leonardob92/LBPD-1.0.git) and the Oxford Centre for Human Brain Activity (OHBA) Software Library (OSL) (available at https://ohba-analysis.github.io/osl-docs/) [30]. OSL is freely available software that builds on the Fieldtrip [31], FSL [32], and SPM [33] toolboxes. Next the continuous MEG data was visually inspected and large artefacts were removed. This removal discarded less than 0.1% of the data. Independent component analyses (ICA) were used to removed artefacts stemming from heart-beats and eye-blinks [34]. Firstly, the original signal was decomposed into independent components. Next, we isolated and discarded the components that picked up activity from eye-blinks and heartbeats. Then the signal was rebuilt using the remaining components. Lastly, the signal was epoched into 135 trials, 27 trials in 5 experimental conditions, and the mean baseline signal, obtained from the post-stimulus brain signal, was removed. Each resulting trial lasted 4500 ms, made up of 4400 ms plus 100 ms of baseline time.

#### D. Source reconstruction

Whilst MEG recordings have excellent temporal resolution when compared to other imaging modalities, one must employ source-reconstruction to spatially locate activity in the brain. We employed the beam-forming algorithm [35–37] implemented in both in-house codes and OSL, SPM and FieldTrip.

In the following, we give a thorough description of the inverse model employed in the beam-forming algorithm. The algorithm is made up of two steps: (1) designing a forward model, (2) computing the inverse solution.

The forward model is a theoretical model that considers each brain source as a voxel/active source. The model describes how the strength of each dipole would be reflected onto each of the MEG sensors. We employed magnetometer channels and an 8-mm grid, which returned 3559 dipole locations (voxels) within the whole brain. We co-registered the individual, structural T1 data with the fiducial points and then computed a forward model by adopting the ‘Single Shelf’ method [38]. This outputs the so-called ‘leadfield’ model which is an *S × M* matrix, *L*, where *S* is the number of sources and *M* is the number of MEG channels. In three cases, the structural T1 was not available and so we performed the leadfield computation with the ‘MNI152-T1 with 8-mm spatial resolution’ template.

Next, we used the beam-forming algorithm to compute the inverse solution. By sequentially applying a set of weights to the source locations, the algorithm can isolate the contribution of each source to the activity recorded by each MEG channel at each time-point of the recording. We summarise the beamforming algorithm in the following steps.

Firstly, the data recorded by the MEG sensors *B* at time *t* is described by the equation,

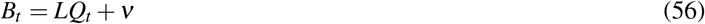

where *L* is the leadfield model, *Q* is the ‘dipole matrix’ which carries the activity of each dipole over time and *ν* is noise [36]. The aim is to compute *Q* by solving the inverse problem. In the beam-forming algorithm, weights are computed and then applied to *B*_*t*_ i.e. for a single dipole, *q*, we have,

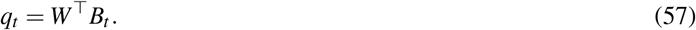

Beam-forming computes weights, *W*_*n*_, for each brain source *n* using the covariance matrix of the MEG sensors, *C*, calculated on the continuous signal with all trials concatenated, in the following fashion,

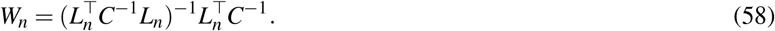

Following Nolte [38], the computation of the leadfield method was performed for three orientations of each brain source. Using singular value decomposition (SVD), the three orientations were reduced to one,

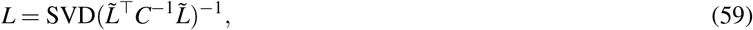

a common technique for simplifying beam-forming output [39, 40]. Here, 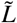 represents the leadfield model with three orientations. Lastly, the obtained weights were applied to each brain source at each time-point and normalised according to Luckhoo et al [40]. In addition to individual trials, the weights were applied to averaged neural activity over all trials. The procedure returned a time-series for each of the 3559 brain sources for each trial, referred to as the ‘neural activity index’. The sign ambiguity of the evoked responses time series was adjusted for each brain source using its sign in correspondence with the N100 response to the first tone of the auditory sequences (see Refs. [23, 25–27]).

Finally, the 3559 voxels obtained through source reconstruction were reduced to six functional brain parcels (or regions of interest (ROIs)) that roughly correspond to auditory cortices in the left and right hemispheres (ACL, ACR); the hippocampal and inferior temporal cortices in the left and right hemispheres (HITL, HITR) and two medial regions, the bilateral medial cingulate gyrus (MC) and the bilateral ventro-medial prefrontal cortex (VMPFC). Our ROIs were taken from the prior study which used this dataset to reveal the event-related neural responses underlying auditory recognition [23]. In said study, an extensive description and validation of these functionally derived ROIs is provided. Moreover, the study showed the consistency of the results when comparing these ROIs with ROIs taken from the well-known Automated Anatomical Labelling (AAL) parcellation and encompassing bilateral Heschl’s gyri, bilateral hippocampi and medial and anterior cingulate cortex. The data was analysed at a temporal resolution of 4 ms so the resulting multivariate time-series was of dimension 6 *×* 1026 (variables *×* time-points).

### XI. SIGNAL QUALITY AND SIGNAL TO NOISE RATIO IN MEG RECORDINGS

In this experimental paradigm, 135 trials were performed per participant for 83 participants, corresponding to 27 trials and 5 experimental conditions [23]. In this study, we focused on a single experimental condition. Furthermore, as analysis is restricted to participants who correctly indicated that the piece of music had not been altered, this means that each participant had a unique number of successful, usable trials up to a maximum of 27. We considered data from the 51 participants who had at least 15 successful, usable trials out of the total 83 participants. For those participants who had more than 15 successful trials, we then randomly sampled 15 trials in order to have an equal contribution from each participant, allowing for the participant level analysis presented in Sec. II. Recommendations for trial counts in an MEG paradigm differ by task, but 40-100 trials is often considered a sensible range [41]. In order to validate our choice to use 15 trials, even for participants who had a larger number of available trials, we consider the participant-trial-averaged signal shown in panel a) of Figure 13. The 6 panels show the average signal of each region, indicated by the schematic representation. The blue lines represent the data used in this study, taken by first averaging over the 15 trials per participant and then averaging over all participants. The shading represents the standard error of each participant compared to the mean signal. The yellow dashed lines represent the mean signal using the total available data with a custom number of trials per participant, with a minimum of 15 and a maximum of 27. This figure highlights the strong neural response associated with this task-paradigm, as found in previous studies [23–28]. Further, the strong similarity between the yellow and blue signals indicates that opting for a smaller number of trials in order to have the same contribution from each participant, does not result in a decrease in signal quality or signal to noise ratio.

In panel b) of Figure 13, we extend this analysis by focusing on the pre-stimulus time, where the red line indicates the stimulus-time. Panel b) shows that the recordings contain a ‘clean’ baseline level of activity with small fluctuations around 0, followed by a strong response after the stimulus.

Each recording contains an epoch of 1026 time-points representing 0.1 seconds of baseline time, followed by a 4 second of active period. The full epoch was considered in the analysis. For further discussion of the paradigm and the associated data see Ref. [23].

### XII. EPOCH COMPARISON

In both the main manuscript and the SI, in order to use the maximum amount of data, we analyse the full epoch of 4.1 seconds, made up of 0.1 seconds of pre-stimulus baseline time and the following 4 seconds of post-stimulus activity. Alternative choices for a suitable epoch are to remove the pre-stimulus baseline of 0.1 seconds and/or to remove the final few seconds of activity where it appears the activity returns to baseline [23]. In order to validate that our results hold irrespective of this choice of epoch, we consider the results of the method applied to three different epochs:

1. **Full epoch:** This is the epoch considered in the main manuscript and the SI (bar this section), which includes 4.1 seconds of activity including 0.1 seconds of pre-stimulus baseline.
2. **Baseline removed:** This epoch contains only the 4 seconds of activity post-stimulus i.e. with the baseline removed.
3. **Baseline removed / Reset removed:** This epoch contains 2.5 seconds of activity, from time 0 to 2.5 i.e. with both the baseline and the subsequent reset to baseline removed.

Figure 14 shows the 4-order irreversibilities applied to the three epoch choices. Panel a) shows that using a shorter epoch results in elevated irreversibility measurements, but that the *relative* behaviour remains the same. Furthermore, the correlations between the results for each epoch are almost perfect with *r* = 1.00 (3 s.f, ****) between Full and Baseline removed, *r* = 0.927 (3 s.f, ****) between Full and Baseline removed/Reset removed and *r* = 0.926 (3 s.f, ****) between Baseline removed and Baseline removed/Reset removed. Therefore the choice of any sensible epoch has minimal impact on the relative results.

---

**FIG. 5.**
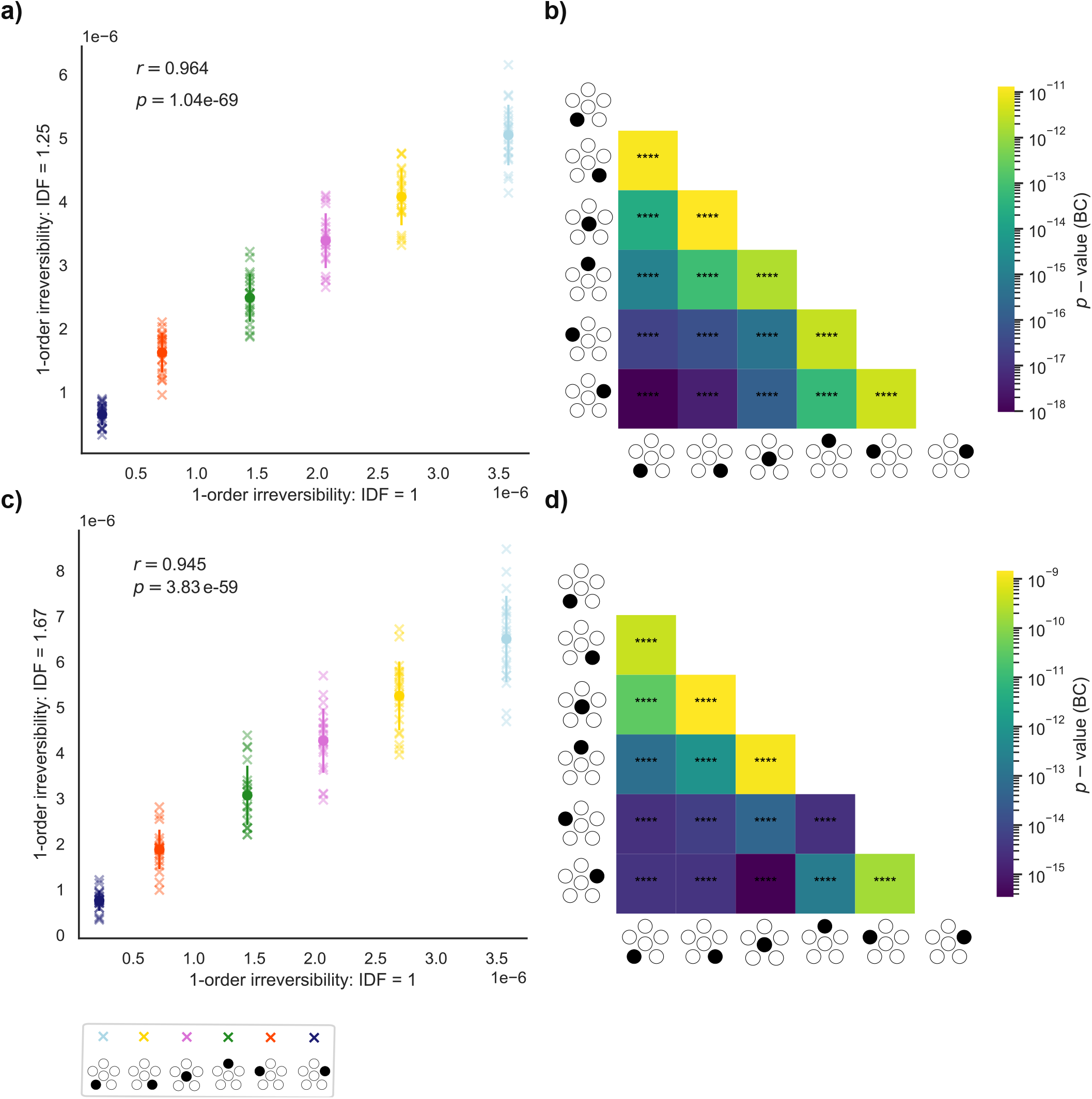
Sub-sampling for estimation of finite-data errors at order 1. a) A comparison between the original results at IDF=1 and the results at 20 samples at IDF=1.25 (12/15 trials). We find that the results are highly correlated indicating that our method is robust to different amounts of data. b) Bonferroni-corrected paired *t*−tests show that each pairwise comparison is significant (****,*p <* 0.0001), indicating that the differences in irreversibilities are far beyond noise-level. c) A comparison between the original results at IDF=1 and the results at 20 samples at IDF=1.67 (9/15 trials). We find that the results are highly correlated. d) Bonferroni-corrected paired *t*−tests show that each pairwise comparison is significant (****,*p <* 0.0001).

**FIG. 6.**
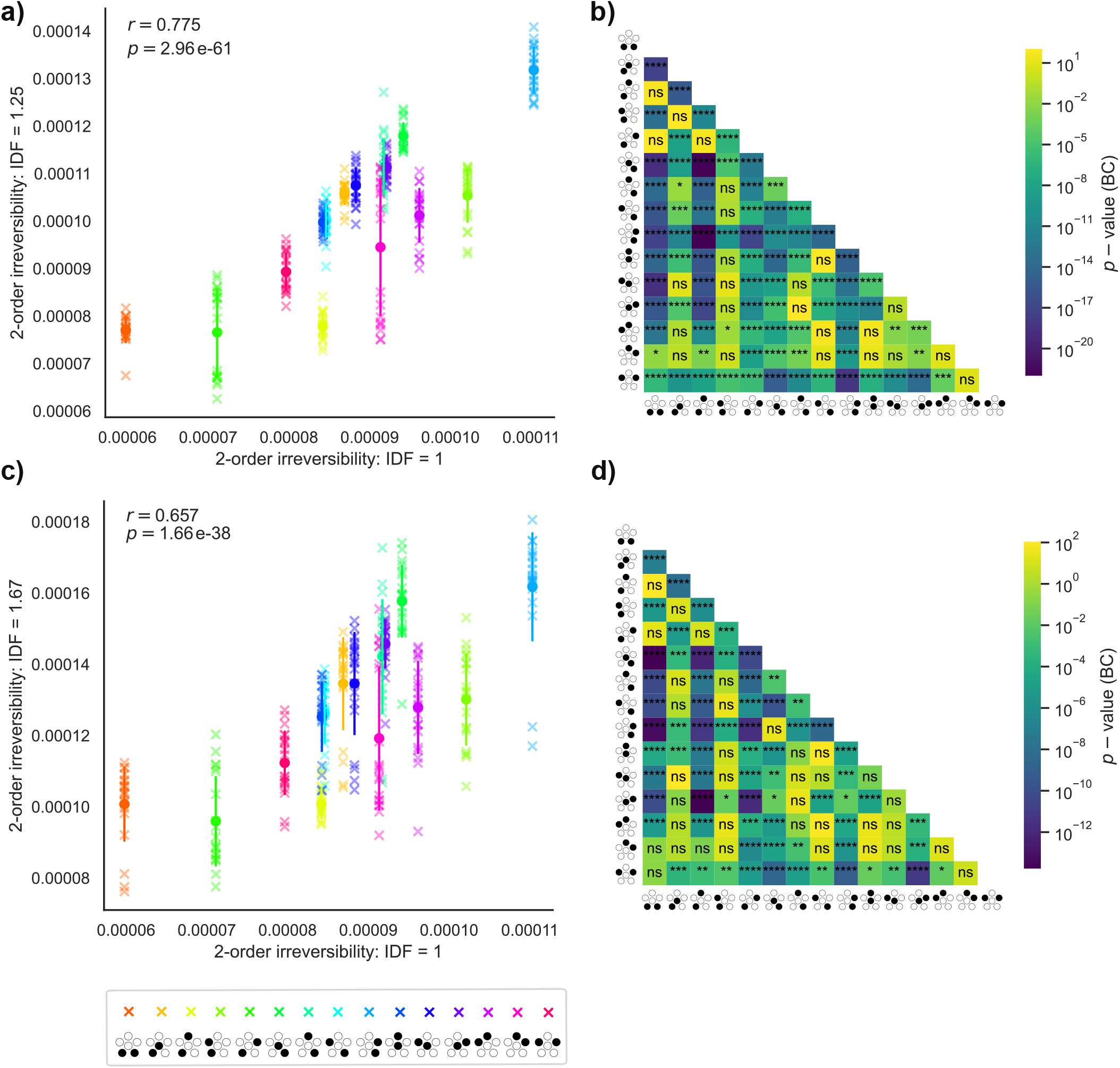
Sub-sampling for estimation of finite-data errors at order 2. a) A comparison between the original results at IDF=1 and the results at 20 samples at IDF=1.25 (12/15 trials). We find that the results are highly correlated. b) Bonferroni-corrected paired *t*−tests show a range of significant pairwise comparisons. c) A comparison between the original results at IDF=1 and the results at 20 samples at IDF=1.67 (9/15 trials). We find that the results are highly correlated. d) Bonferroni-corrected paired *t*−tests show a range of significant pairwise comparisons.

**FIG. 7.**
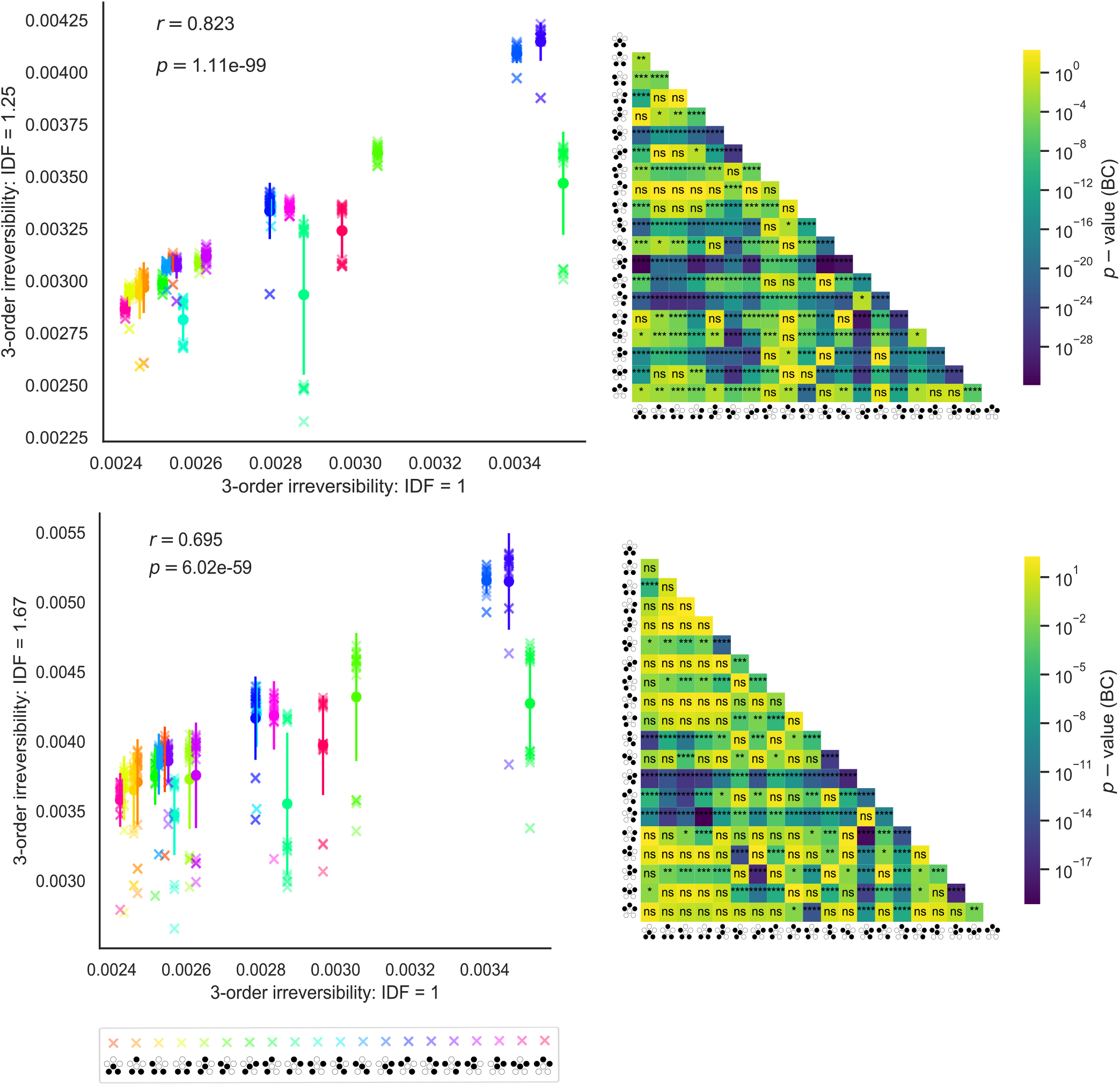
Sub-sampling for estimation of finite-data errors at order 3. a) A comparison between the original results at IDF=1 and the results at 20 samples at IDF=1.25 (12/15 trials). We find that the results are highly correlated. b) Bonferroni-corrected paired *t*−tests show a range of significant pairwise comparisons. c) A comparison between the original results at IDF=1 and the results at 20 samples at IDF=1.67 (9/15 trials). We find that the results are highly correlated. d) Bonferroni-corrected paired *t*−tests show a range of significant pairwise comparisons.

**FIG. 8.**
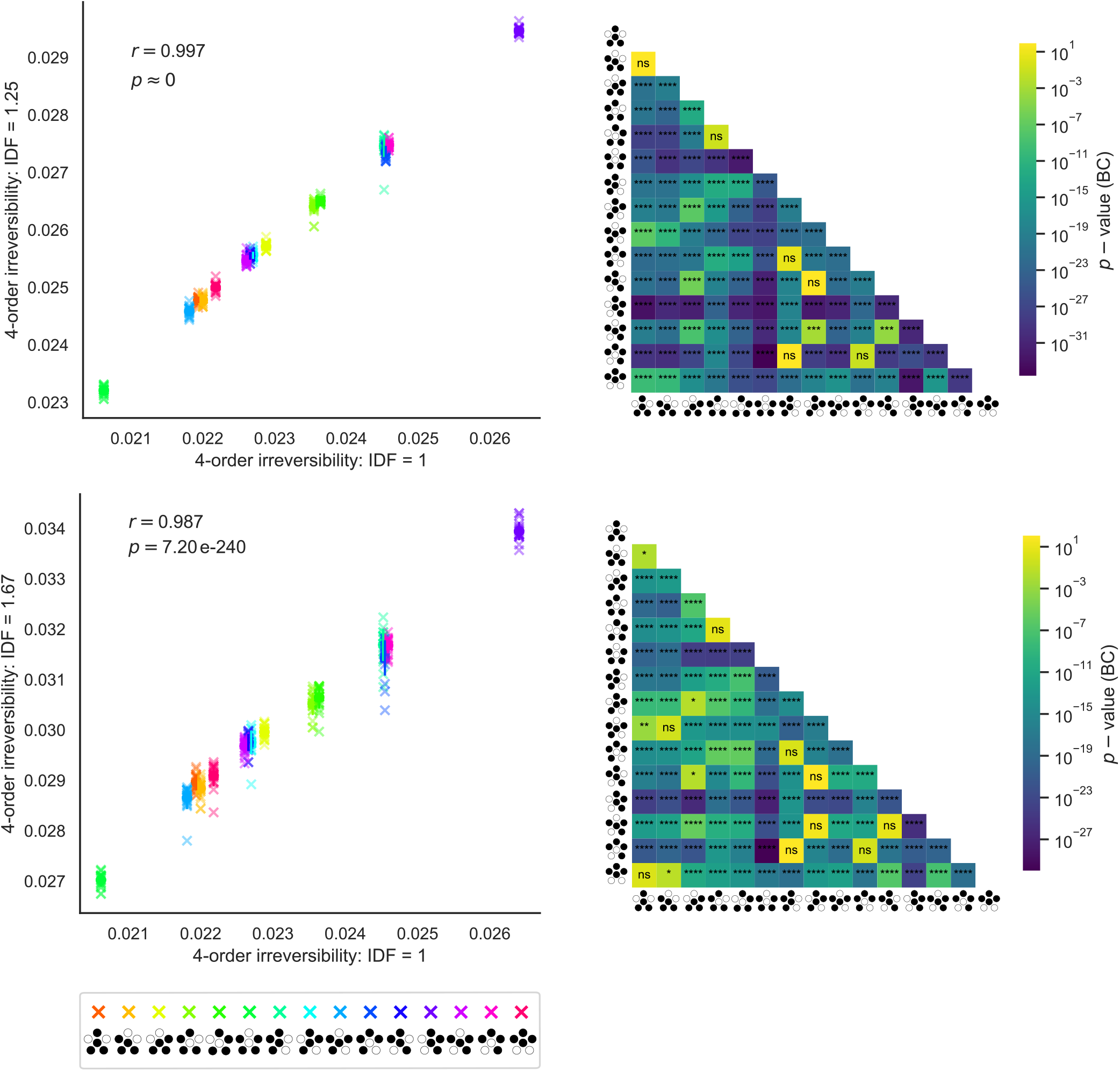
Sub-sampling for estimation of finite-data errors at order 4. a) A comparison between the original results at IDF=1 and the results at 20 samples at IDF=1.25 (12/15 trials). We find that the results are highly correlated. b) Bonferroni-corrected paired *t*−tests show a range of significant pairwise comparisons. c) A comparison between the original results at IDF=1 and the results at 20 samples at IDF=1.67 (9/15 trials). We find that the results are highly correlated. d) Bonferroni-corrected paired *t*−tests show a range of significant pairwise comparisons.

**FIG. 9.**
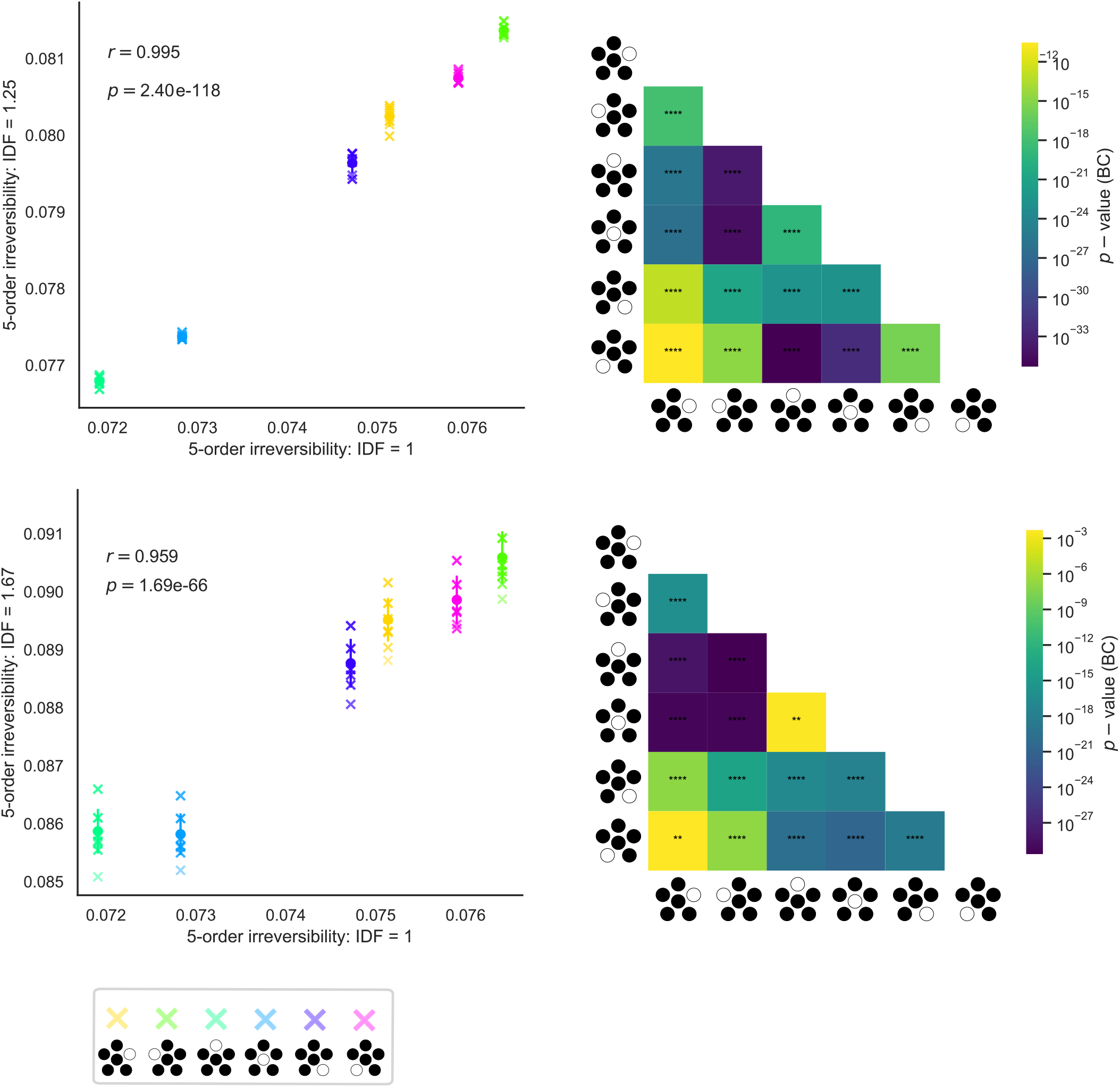
Sub-sampling for estimation of finite-data errors at order 5. a) A comparison between the original results at IDF=1 and the results at 20 samples at IDF=1.25 (12/15 trials). We find that the results are highly correlated. b) Bonferroni-corrected paired *t*−tests show a range of significant pairwise comparisons. c) A comparison between the original results at IDF=1 and the results at 20 samples at IDF=1.67 (9/15 trials). We find that the results are highly correlated. d) Bonferroni-corrected paired *t*−tests show a range of significant pairwise comparisons.

**FIG. 10.**
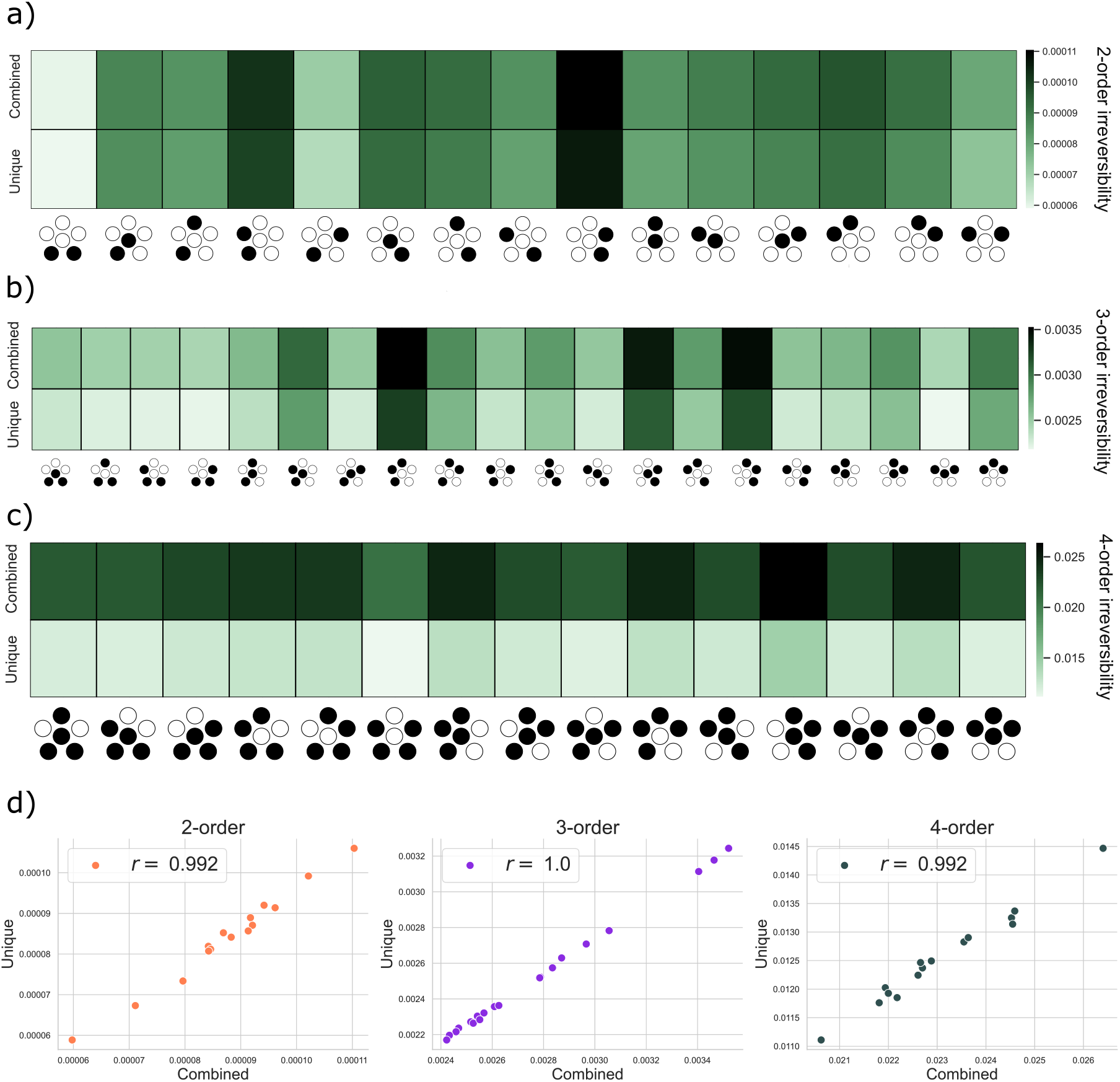
Comparison of the combined *k*-order contributions to irreversibility against the unique contributions from each *k*-tuple for *k* = 2, 3, 4. a) *k* = 2. As the irreversibility of the pairwise interactions is much larger than the individual trajectories, the unique and combined irreversibilities are very similar. b) *k* = 3. Whilst there is some contrast between the unique and combined irreversibilities, the general hierarchy is preserved. c) *k* = 4. Again there is some contrast between the unique and combined irreversibilities with the general hierarchy being preserved. d) We show the almost perfect correlation between the unique and combined irreversibilities at each level. This indicates that the *k*−order interactions dominate the irreversibility at level *k* and suggest little difference when considering unique or combined irreversibilities.

**FIG. 11.**
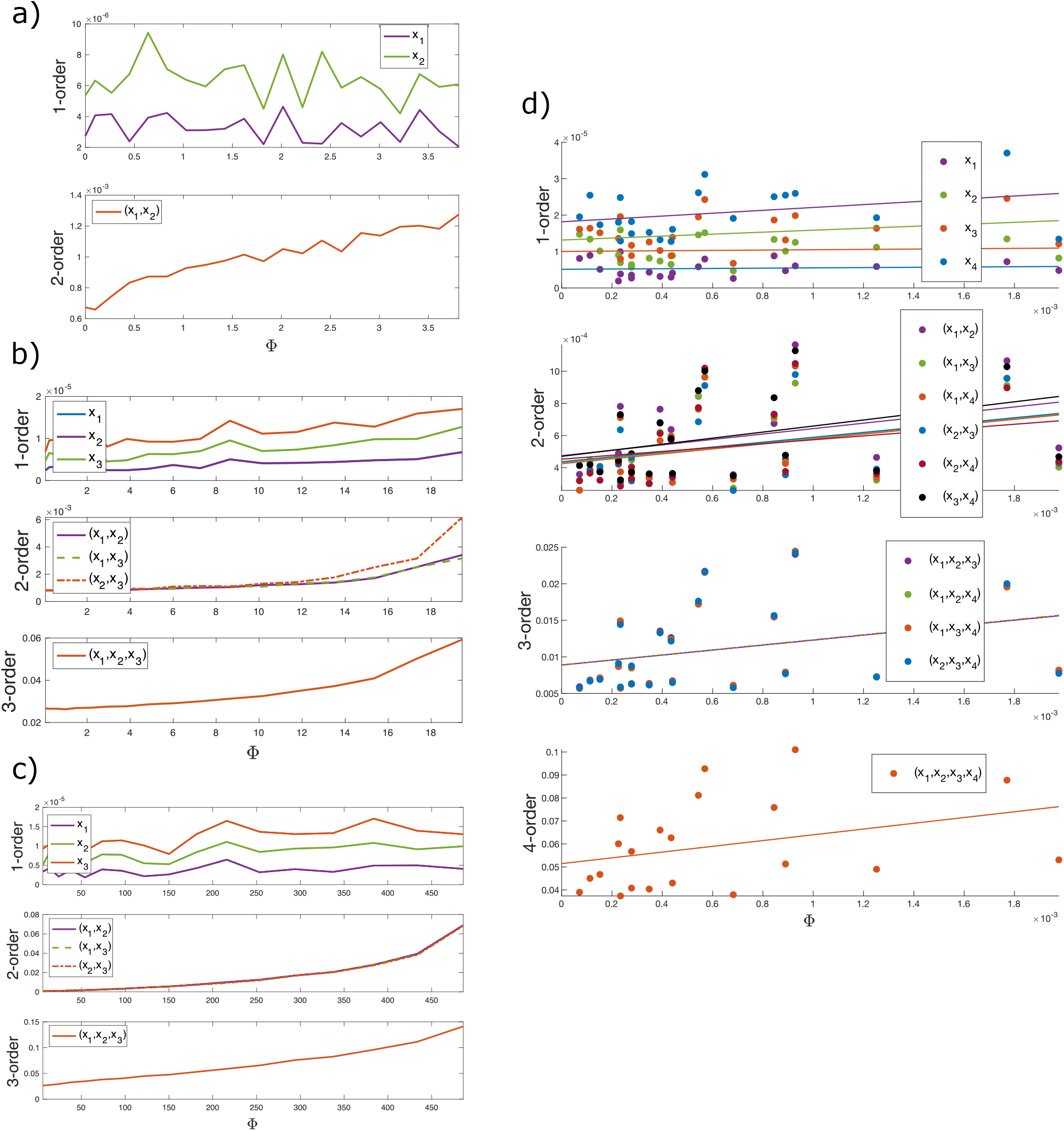
Validation of the DiMViGI framework using simulated data from the mOU. a) Example 1 - a 2-dimensional process. The pairwise irreversibility scales with the global rate Φ, whilst the individual variables do not. b) Example 2 - a 3 dimensional process with drift-disjoint pairs. The pairs and triplet irreversibilities scale with the global rate Φ whilst the individual trajectories do not. c) Example 3 - a 3 dimensional process with 3 way interactions in the drift and noise. The pairs and triplet irreversibilities scale with the global rate Φ whilst the individual trajectories do not. d) Example 4 - a 4 dimensional process with 2 strongly interacting pairs. The global rate Φ is estimated numerically producing variance in the plot so we plot least-square regressions. The pairs and triplets and quadruplet irreversibilities scale with the global rate Φ whilst the individual trajectories do not. Notably, the strongly interacting pairs (*x*_1_, *x*_2_) and (*x*_3_, *x*_4_) have a higher level of irreversibilities than the other pairs.

**FIG. 12.**
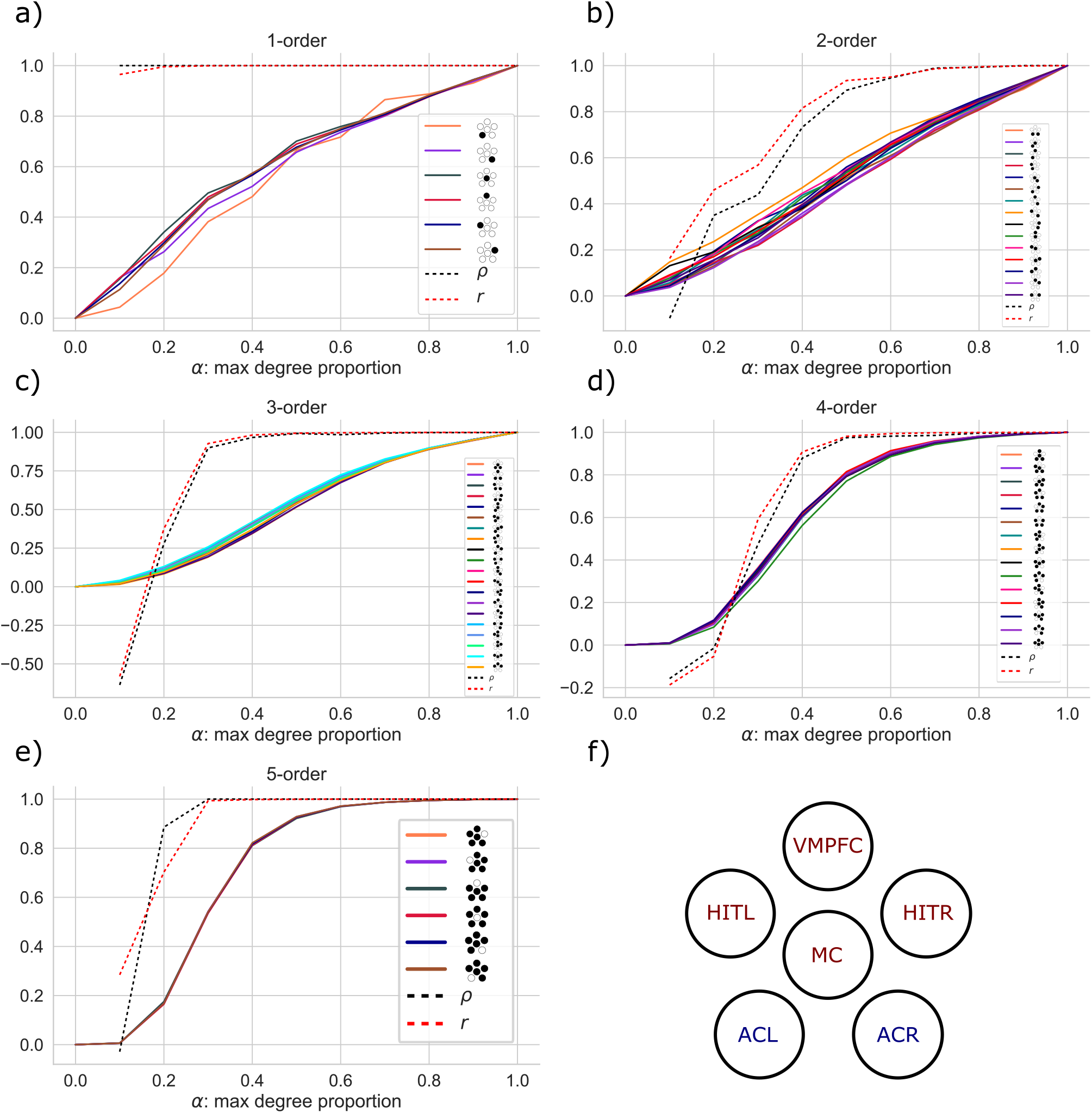
Systematic analysis of the effect of degree-limiting on the irreversibility of each tuple. Panels a-e) show the proportion of irreversibility captured with degree limited to 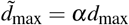 for *α* ∈ [0, 1]. Using a limited degree underestimates the irreversibility of the tuple. In addition a-e) show the correlation between the limited degree results at each level, and the full degree results. For higher orders, degree limiting is shown to lose minimal information, both in terms of the correlations between tuples and absolute values. This indicates that is is a valuable technique for maximising the efficiency of the DiMViGI framework. Panel f) recalls the schematic representation for the tuples in the legends.

**FIG. 13.**
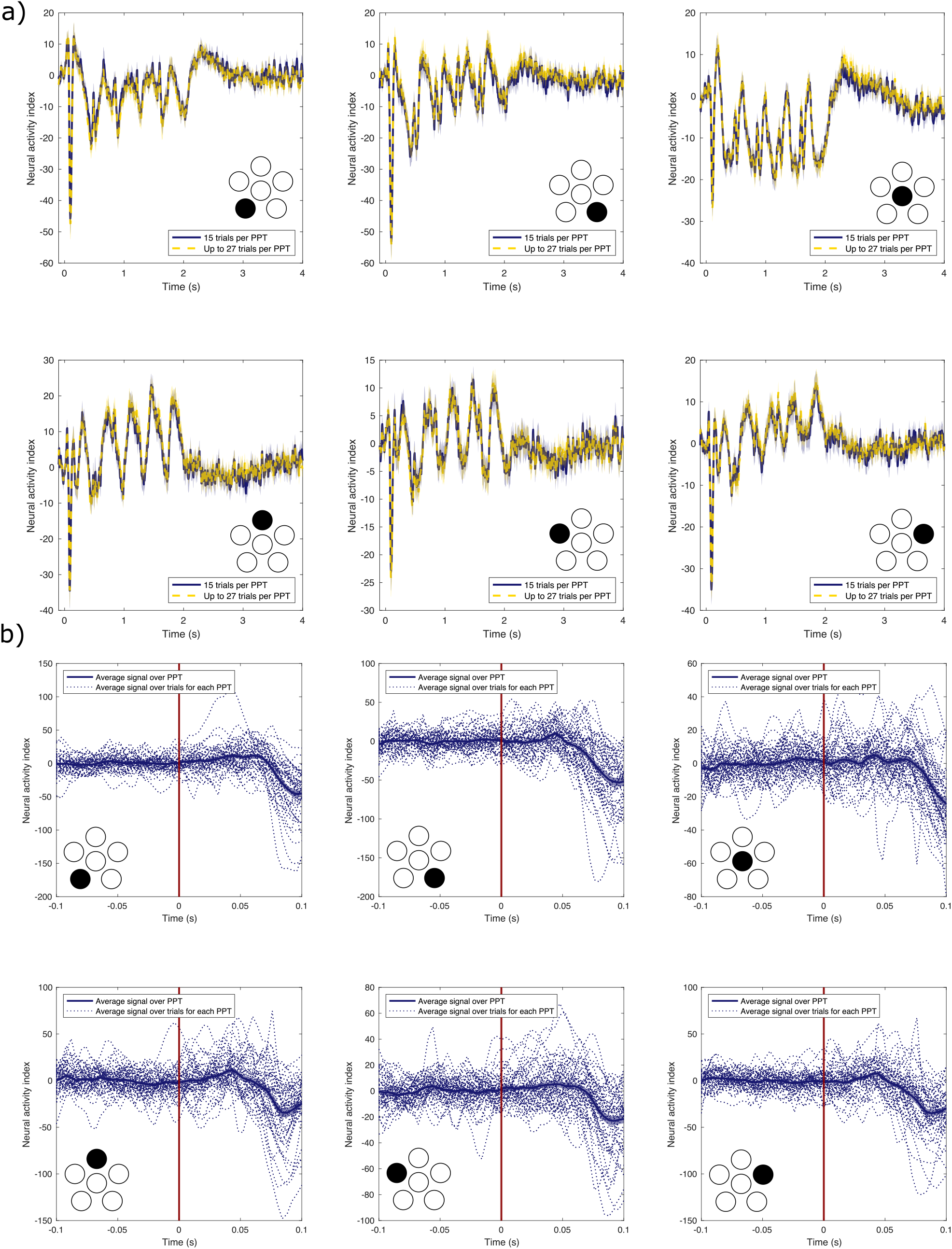
Signal visualisation. a) The 6 panels represent the averaged signal for each of the ROIs, represented by the schematic diagram. The blue lines represent the averaged signal of the data used in this analysis. This is obtained by averaging the signal for each participant over their associated 15 trials. Next, we average over all participants, measuring the standard error for each participant compared to the mean. The red line contains the same analysis but using all possible trials for each participant, between 15 and 27 trials. b) Each pane represents the signal of each participant as well as the mean, of the data used in this study, focused on the baseline, prestimulus time. The red line denotes the stimulus time. The small random fluctuations before the stimulus time indicate a ‘clean’ baseline.

**FIG. 14.**
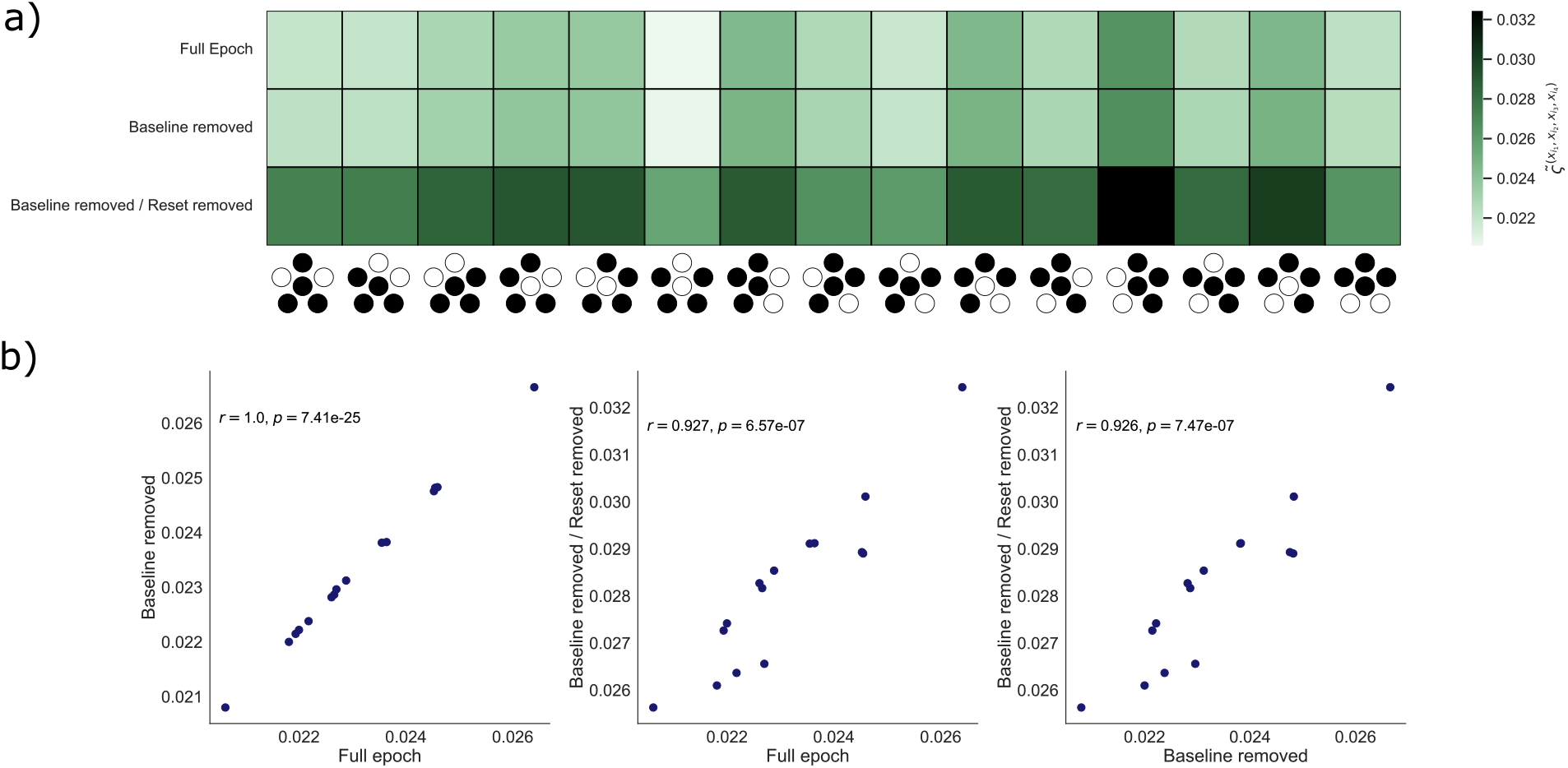
Epoch comparison. a) The results of the 4-order irreversibilities for each of the three choices of epoch yield very similar results. The overall level of irreversibility is elevated for the shortest epoch. b) The correlations between the results for each epoch length are almost perfect suggesting that each choice yields equivalent results.

## Notes

### Competing Interest Statement

The authors have declared no competing interest.

### Summary of Updates

Our revised manuscript includes a large number of additional analyses for validating the quality of our experimental data and the rigour and robustness of our analytical approach. These include, but are not limited to: 1. Visualisations of the pre-processed MEG recordings to showcase the strong neural response associated with our experimental paradigm, as well as to validate a clean baseline and a strong signal-to-noise-ratio. 2. An example comparison of the results obtained with epochs of different duration. 3. Employment of a sub-sampling approach to estimate the size of finite-data errors and validate the significance of difference between the irreversibility measured in different tuples. In addition, we have made many changes to the written content of the manuscript that improve the clarity and rigour of our methods and results, as well as broader discussion of our paper within the wider literature of a number of inter-disciplinary fields including the neuroscience of the auditory system, network neuroscience, higher-order interactions and non-equilibrium dynamics.

